# Calcium signals shape metabolic control of H3K27ac and H3K18la to regulate EGA

**DOI:** 10.1101/2025.03.14.643362

**Authors:** Virginia Savy, Paula Stein, Don Delker, Martín A. Estermann, Brian N. Papas, Zongli Xu, Lenka Radonova, Carmen J. Williams

## Abstract

The use of assisted reproductive technologies (ART) has enabled the birth of over 9 million babies; but it is associated with increased risks of negative metabolic outcomes in offspring. Yet, the underlying mechanism remains unknown. Calcium (Ca^2+^) signals, which initiate embryo development at fertilization, are frequently disrupted in human ART. In mice, abnormal Ca^2+^ signals at fertilization impair embryo development and adult offspring metabolism. Changes in intracellular Ca^2+^ drive mitochondrial activity and production of metabolites used by the epigenetic machinery. For example, acetyl-CoA (derived mainly from pyruvate) and lactyl-CoA (derived from lactate) are used for writing H3K27ac and H3K18la marks that orchestrate initiation of development. Using both a genetic mouse model and treatment with ionomycin to raise intracellular Ca^2+^ of wild-type fertilized eggs, we found that excess Ca^2+^ at fertilization changes metabolic substrate availability, causing epigenetic changes that impact embryo development and offspring health. Specifically, increased Ca^2+^ exposure at fertilization led to increased H3K27ac levels and decreased H3K18la levels at the 1-cell (1C) stage, that persisted until the 2-cell (2C) stage. Ultralow input CUT&Tag revealed significant differences in H3K27ac and H3K18la genomic profiles between control and ionomycin groups. In addition, increased Ca^2+^ exposure resulted in a marked reduction in global transcription at the 1C stage that persisted through the 2C stage due to diminished activity of RNA polymerase I. Excess Ca^2+^ following fertilization increased pyruvate dehydrogenase activity (enzyme that converts pyruvate to acetyl-CoA) and decreased total lactate levels. Provision of exogenous lactyl-CoA before ionomycin treatment restored H3K18la levels at the 1C and 2C stages and rescued global transcription to control levels. Our findings demonstrate conclusively that Ca^2+^ dynamics drive metabolic regulation of epigenetic reprogramming at fertilization and alter EGA.

## Introduction

In mammals, development of the newly formed embryo is initiated by sperm-induced oscillatory calcium (Ca^2+^) signals^1,2^. These signals, which last only for the first few hours after fertilization, trigger a series of critical transitions required for successful onset and progression of preimplantation embryo development^3,4^. Disruption of the Ca^2+^ signals impairs the competence of mouse embryos to complete development to term and results in sexually dimorphic alterations in offspring growth and body composition^5–7^. It is unknown how Ca^2+^ signals are mechanistically connected to these long-term metabolic changes. This information is critical because assisted reproductive technologies (ART) can dramatically modify fertilization induced Ca^2+^ dynamics^8–10^. In fact, Ca^2+^ ionophores are currently used in human clinics to artificially raise Ca^2+^ levels after in vitro fertilization (IVF) as a treatment for recurrent fertility failure, despite being poorly researched and carrying long-term consequences^11,12^. Moreover, cellular Ca^2+^ homeostasis is disrupted in various tissues of patients with medical conditions like obesity, diabetes and inflammation, likely affecting female gametes and their ability to handle Ca^2+^ signals at fertilization^13^.

Upon fertilization, as the egg’s Ca^2+^ levels oscillate, there is a rapid global rearrangement of epigenetic modifications essential for successful development^14^. Two of these are post-translational modifications of histone H3: H3 lysine 27 acetylation (H3K27ac; associated with open chromatin) and H3 lysine 18 lactylation (H3K18la; enriched at active enhancers)^15–17^. Both H3K27ac and H3K18la are enriched in the maternal and paternal pronuclei, promoting an overall open chromatin state that facilitates the initial phase of embryonic genome activation (EGA), known as minor EGA^18–20^. In mice, minor EGA takes place at the late 1-cell (1C) stage, is generally promiscuous and includes both RNA polymerase I (Pol-I)-mediated transcription of ribosomal DNA and RNA polymerase II (Pol-II)-mediated transcription of mRNA^21,22^. As development proceeds through the 2-cell (2C) stage, there is a global loss of H3K27ac while H3K18la remains relatively high, and chromatin accessibility becomes more restricted^14,18^. By the late 2C stage, when the large burst of transcriptional activity known as major EGA begins, promiscuous transcription has ended and genes required for the 2C stage and beyond are transcribed^23^. The rapid, dynamic, yet carefully regulated shift in epigenetic states during this brief developmental window is critical for successful EGA and progression of embryonic development.

Metabolism plays a crucial role in regulating chromatin structure by controlling metabolite levels that serve as cofactors for the epigenetic machinery. For example, histone acetyltransferases write H3K27ac marks using acetyl-CoA, which is in part generated from pyruvate by the mitochondrial pyruvate dehydrogenase (PDH) complex^24^. Sirtuin 1 (SIRT1), the deacetylase responsible for removing H3K27ac marks in 2C embryos, is regulated by metabolic changes in NAD+:NADH ratios^18^. In fact, lack of NAD+ during the 1C-2C transition leads to developmental arrest at the 2C stage, with excessive H3K27ac and aberrant expression of 1C-related genes that should normally be silenced by this time. A growing body of evidence points toward lactate levels and protein lactylation as a molecular switch bridging metabolic changes and epigenetic reprogramming^25,26^. Lactate is enriched in the nucleus of 2C embryos, and lactate dehydrogenase (LDH) activity is crucial for EGA in mice and humans due to its role in providing lactyl-CoA for writing H3K18la^19,27^. Similarly, a metabolic switch involving lactate levels and protein lactylation regulates the transition between embryonic stem cells and 2-cell-like cells^28^. These examples highlight the interconnections between metabolic shifts and epigenetic reprogramming that are important in both somatic cells and early embryos.

Cytosolic Ca^2+^ levels act as a rapid and sensitive switch regulating metabolism. Ca^2+^ allosterically boosts the activity of mitochondrial enzymes, including PDH, that drive the tricarboxylic acid (TCA) cycle, stimulating production of NADH and ATP^29^. In fertilized mouse eggs, cytosolic Ca^2+^ oscillations cause simultaneous oscillations in mitochondrial Ca^2+^ that drive ATP production required to maintain the cytosolic Ca²^+^ oscillations and overall cellular Ca^2+^ homeostasis^30,31^. Interestingly, several TCA cycle enzymes, including PDH, migrate from the mitochondria to the nucleus in early mouse embryos and in induced pluripotent stem cells, presumably to provide local metabolites for chromatin remodeling during EGA and totipotency acquisition^32,33^. Here we used an orthogonal experimental design to identify the link between Ca^2+^ homeostasis, metabolic status, and epigenetic reprogramming during the maternal-to-zygotic transition. Using both a mouse knockout model and ionomycin treatment of early embryos we show that abnormal Ca²^+^ exposure following fertilization impacts metabolism and redox status, resulting in changes in H3K27ac and H3K18la, and impaired minor EGA and preimplantation development. Adult offspring derived from the knockout females have altered metabolism and differences in DNA methylation of genes that regulate visceral fat. These data provide an underlying mechanism for the “butterfly effect” that post-fertilization Ca^2+^ signals have on adult metabolic set points.

## Results

### 1. Ca²^+^ responses are dramatically disrupted in fertilized PMCA1/PMCA3-null eggs, impacting their metabolic status

To investigate mechanisms linking changes in Ca^2+^ signaling at the 1C stage with offspring metabolism, we developed a mouse model of dramatically increased Ca^2+^ exposure at fertilization in vivo. *Atp2b3*, which encodes the plasma membrane Ca^2+^ ATPase 3 (PMCA3), was globally deleted in zygotes that already had a floxed (f) allele of *Atp2b1* (encodes PMCA1)^5^. Conditional deletion of *Atp2b1* in oocytes was accomplished using the Zp3-cre transgene^34^. The final experimental mice were females carrying PMCA1/PMCA3-null eggs (dKO), PMCA3-null eggs (3KO), or PMCA1-f/f eggs (Ctrl). Comparison of Ca^2+^ dynamics at fertilization revealed that, while 3KO eggs had a small, ~1.6-fold increase in the length of the first Ca^2+^ transient, dKO eggs had extended elevated Ca^2+^ at fertilization, a 20-fold increase over Ctrl (Fig. 1A, 1B). The area under the Ca^2+^ curve (AUC), representing the accumulated Ca^2+^ exposure that is considered the main driver of embryo development^4^, was not altered in 3KO eggs but was 10-fold higher in dKO eggs relative to Ctrl (Fig. 1C). To account for potential off target effects of gene targeting, two independent mouse sublines were generated and analyzed. The Ca^2+^ phenotypes in the two dKO sublines were indistinguishable (Fig. S1A, S1B), so subsequent analyses used only one line. The elevated Ca^2+^ response was not due to increased levels of endogenously stored Ca^2+^, suggesting that the higher Ca^2+^ exposure results from impaired Ca^2+^ clearance due to the absence of PMCAs (Fig. S1C).

**Figure 1.**
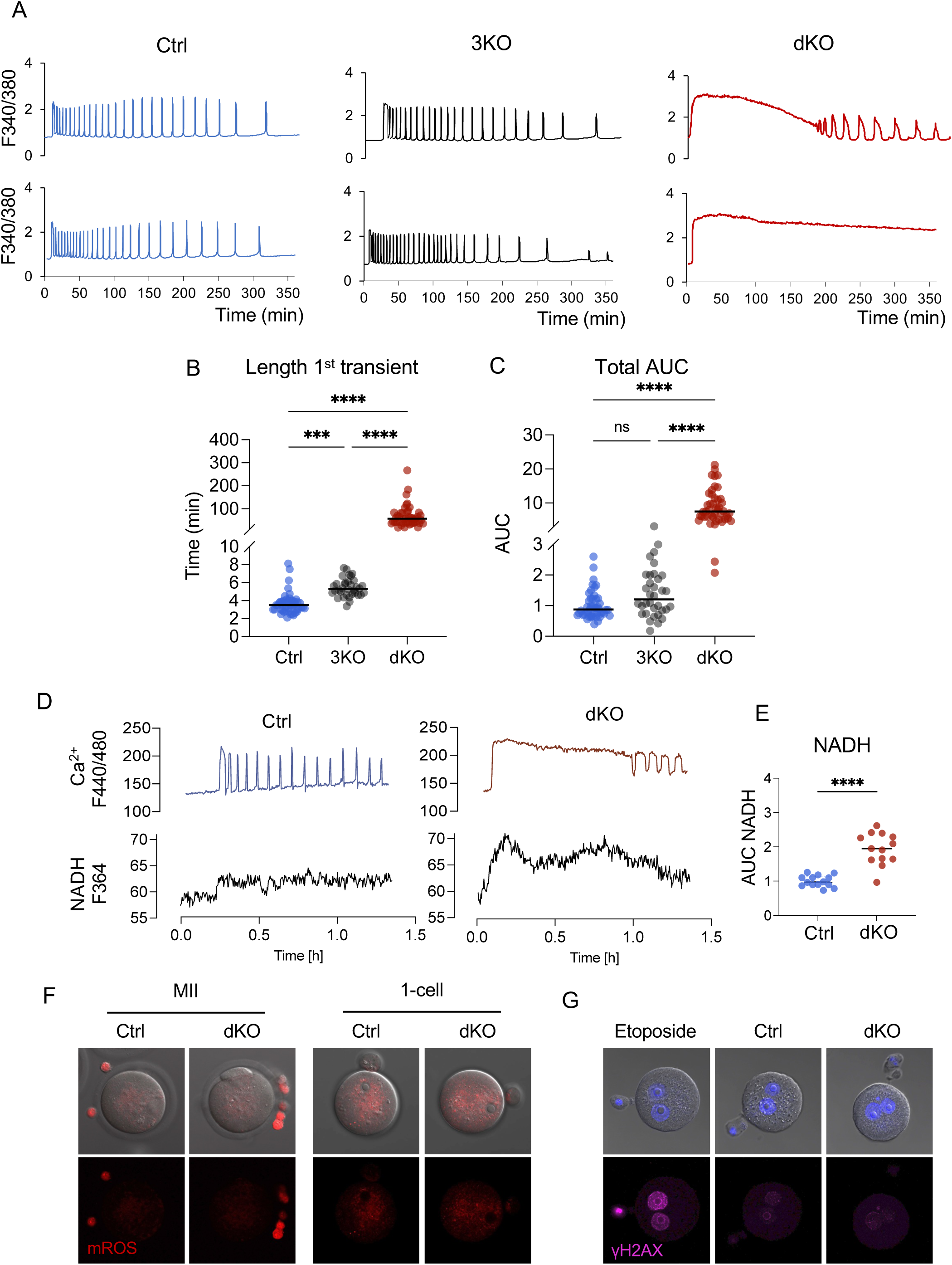
Ca^2+^ responses are dramatically disrupted in PMCA1/PMCA3-null eggs. Ratiometric Ca^2+^ imaging during IVF of control (Ctrl, blue), PMCA3-KO (3KO, gray) and PMCA1/PMCA3 double KO (dKO, red) eggs. N = 3 independent experiments; (A) representative Ca^2+^ traces. (B) Length of the first Ca^2+^ transient. (C) Area under the curve (AUC) of Ca^2+^ signal, relative to controls. Horizontal bars indicate median. For (A-C) Kruskal-Wallis test with Dunnett’s multiple comparisons test were used; ns, not significant; **** p<0.0001. (D) Representative Ca^2+^ response (top) and NADH production (bottom) of Ctrl and dKO eggs during fertilization. (E) AUC of NADH levels, relative to controls. Unpaired t-test, **** p<0.0001. (F) Representative images of mitochondrial reactive oxygen species (mROS) levels detected with mitoSox red in MII eggs and 1C embryos from control and dKO females. (G) Representative images showing DNA damage levels in 1C embryos, detected using the γH2AX antibody. Samples include embryos from control and dKO females, as well as control embryos cultured with etoposide as a positive control.

Sperm-triggered Ca^2+^ oscillations directly regulate mitochondrial metabolism, enhancing TCA cycle activity to supply NADH, FADH2 and ultimately ATP, which is necessary to sustain the energetically demanding Ca²⁺ oscillations^30,31^. As anticipated, NADH levels increased in parallel with Ca^2+^ levels at fertilization and were abnormally elevated in dKO eggs (Fig. 1D, 1E), suggesting an increased mitochondrial activity in this group. However, increased TCA cycle activity can cause production of cytotoxic reactive oxygen species (ROS) and can be associated with DNA damage^35,36^. We therefore tested the impact of prolonged high Ca^2+^ exposure on ROS levels before and after fertilization, and DNA damage in pronuclear stage embryos. Mitochondrial ROS levels were very low prior to fertilization and similar between Ctrl- and dKO-derived 1C embryos (Fig. 1F). Consistent with normal ROS levels, dKO and Ctrl embryos had comparable γH2AX levels after fertilization (Fig. 1G). These results demonstrate that lack of PMCA1 and PMCA3 dramatically disrupts sperm-induced Ca^2+^ oscillations and alters the metabolic status of the fertilized egg without causing persistent ROS production or DNA damage.

### 2. Females carrying dKO eggs are subfertile due to impaired preimplantation embryo development

We next characterized the impact of increased Ca²⁺ exposure on reproductive outcomes in the 3KO and dKO mouse models. Both 3KO and dKO females yielded same number of ovulated eggs as Ctrl females after hormonal stimulation (Fig. 2A). When females were bred for 6 months to wild type (WT) males, there was no difference in the number of litters delivered (Fig. S2A). Although litter size was comparable between 3KO and Ctrl dams, dKO dams had fewer pups per litter with an average litter size approximately 60% of controls (Fig. 2B). dKO dams consistently delivered smaller litters compared to both 3KO and Ctrl dams, indicating a persistent reduction in fertility rather than ovarian failure with age (Fig. 2C). Because the 3KO eggs did not exhibit excess Ca^2+^ exposure (Fig. 1C) and had normal litter sizes (Fig. 2B), subsequent analyses were focused only on the dKO eggs.

**Figure 2.**
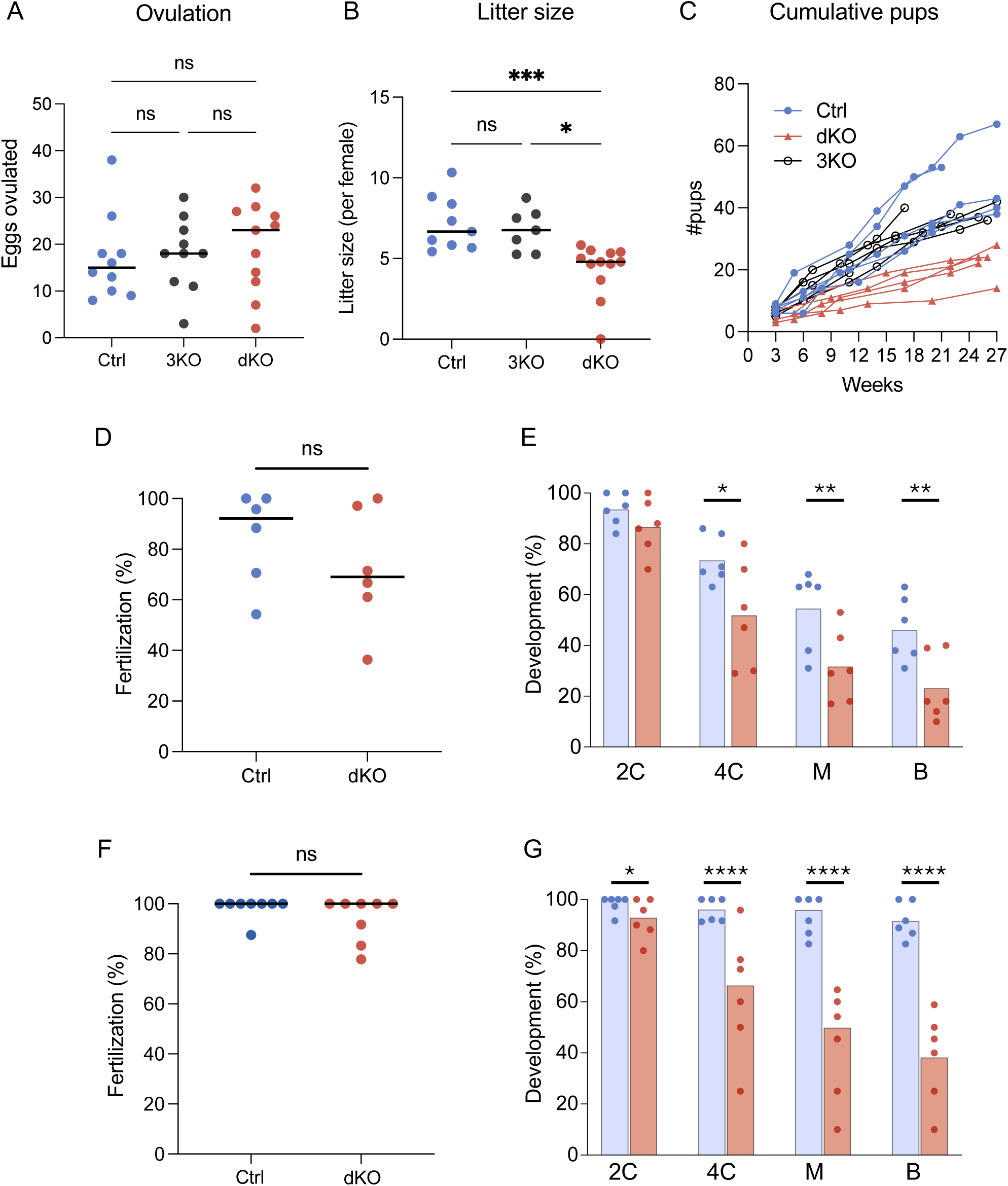
Females carrying dKO eggs are subfertile due to impaired preimplantation embryo development. Fertility assessment of control (Ctrl, Blue), PMCA3-knockout (3KO, grey) and double-KO (dKO, red) females. (A) Number of eggs per superovulated female. (B) Live pups per litter after mating Ctrl, 3KO, and dKO females to WT males; each dot represents one litter from 9, 7 and 11 breeding pairs, respectively. For (A-B) horizontal bars indicate median. Kruskal-Wallis with Dunnett’s multiple comparisons test; *p<0.05; ***p<0.001. (C) Cumulative numbers of viable offspring produced by 5 representative females from each genotype during a 6-month breeding trial. (D-G) Impact of excess Ca^2+^ on fertilization and embryo development. Percentage of fertilized eggs following fertilization in vitro (D) or in vivo (F). Each dot represents an independent biological replicate and horizontal bars indicate median. Mann Whitney test; ns, not significant. In vitro fertilization was performed following zona-pellucida removal. A total of 88 Ctrl and 90 dKO zona-free eggs were included in the analysis. In vivo fertilized eggs were flushed from the oviduct at the 1C stage, and a total of 135 Ctrl and 145 dKO fertilized eggs were included in the analysis. (E, G) Percentage of embryos in panels D and F, respectively, to reach the various preimplantation embryo stages. Each dot represents an independent biological replicate. Preimplantation embryo stages 2C (2-cell), 4C (4-cell), M (morula) and B (blastocyst) stage embryos. For embryo development, the Chi-square test for proportions was used; *p<0.05, **p<0.01, ****p<0.0001.

To determine the timing of the elevated Ca^2+^ impact on fertility, dKO and Ctrl eggs were inseminated in vitro and monitored for preimplantation development. When standard zona pellucida (ZP)-intact IVF was performed, only about 20% of the dKO eggs were fertilized (Fig. S2B). Furthermore, no sperm were visualized in the perivitelline space of the unfertilized eggs, indicating a failure of sperm to penetrate the ZP. This result was unexpected, as natural mating trials showed only a 40% decrease in litter size (Fig. 2B), compared to the 80% reduction in fertilization observed during IVF. These findings suggested that the poor IVF outcome for dKO eggs was a consequence of the in vitro manipulation process. One explanation for the lack of sperm penetration was premature cortical granule release. Under physiological conditions, the first Ca^2+^ rise at fertilization causes cortical granule exocytosis and release of enzymes that modify the ZP, preventing polyspermy^37^. Prior to insemination, a normal pattern of cortical granules was detected in Ctrl eggs but dKO eggs had very few (Fig. S2C), suggesting that premature cortical granule release had occurred. This was indeed the case as demonstrated by premature cleavage of ZP2 protein to the ZP2f form (Fig. S2D). These results suggest that the in vitro manipulation process itself can disrupt Ca^2+^ homeostasis in eggs. In the case of dKO eggs, because they cannot rapidly clear cytosolic Ca^2+^, certain Ca^2+^-triggered events may be prematurely activated, including cortical granule release and cleavage of ZP2 protein. As expected, when the ZP was removed prior to insemination, fertilization was restored to control levels and cleavage to the 2C stage was similar between groups (Fig. 2D-E). However, further development of dKO embryos beyond 2C was impaired, and only half the percentage of dKO as Ctrl embryos reached the blastocyst stage (Fig. 2E), suggesting that abnormal Ca^2+^ homeostasis at fertilization induces changes that persist beyond the 2C stage and impair normal developmental progression.

To confirm the developmental compromise in dKO eggs, eggs fertilized in vivo were recovered from the oviducts of dKO and control dams mated to WT males. There was no difference in the number of fertilized eggs between groups (Fig. 2F), confirming that the low IVF success was a result of the in vitro manipulation. dKO embryos were very slightly compromised in cleaving from 1C to 2C (Fig. 2G). Similar to the ZP-free embryos, only 40% of dKO 1C embryos reached the blastocyst stage compared to 90% of controls. These numbers aligned with the significant reduction (about 40%) in litter size. As observed for the IVF embryos, many of these in vivo-derived dKO embryos arrested their development at the 2C or 4C stages. Together, these results demonstrate that lack of PMCA1 and PMCA3 does not impair oocyte growth or fertilization in vivo and instead impairs the preimplantation developmental program. Moreover, the arrest of embryo development at the 2C-4C stage suggests that excess Ca^2+^ at fertilization negatively affects the maternal-to-zygotic transition.

### 3. Increased Ca^2+^ at fertilization alters offspring metabolic parameters

Moderately increased or decreased Ca^2+^ levels at fertilization are associated with changes in offspring growth trajectory and metabolism^5–7^. To assess long-term effects of dramatically altered Ca^2+^ exposure in the embryos that survived to term, offspring growth, body composition and metabolic parameters of genotype-matched dKO and control-derived pups were analyzed. Offspring from dKO and control dams had comparable weights at birth and gained similar amounts throughout development (Fig. 3A). Although there was a trend toward increased fat content in dKO pups, the difference was not statistically significant in offspring at 5-6 months of age (Fig. 3B). Body mineral composition was comparable between groups, suggesting no effects on bone development (Fig. S3A, S3B). To assess metabolic alterations, animals underwent glucose tolerance testing (GTT). Male offspring derived from dKO mice had much higher serum glucose levels after a 14-h fast, suggesting a persistent impact on metabolism (Fig. 3C). However, the response to the glucose challenge was similar between dKO and control offspring, as indicated by comparable AUC values (Fig. 3D). These results from our dKO mouse model closely resemble observations for IVF-conceived children, who exhibit higher fasting glucose levels but maintain a normal response to metabolic challenges^38,39^.

**Figure 3.**
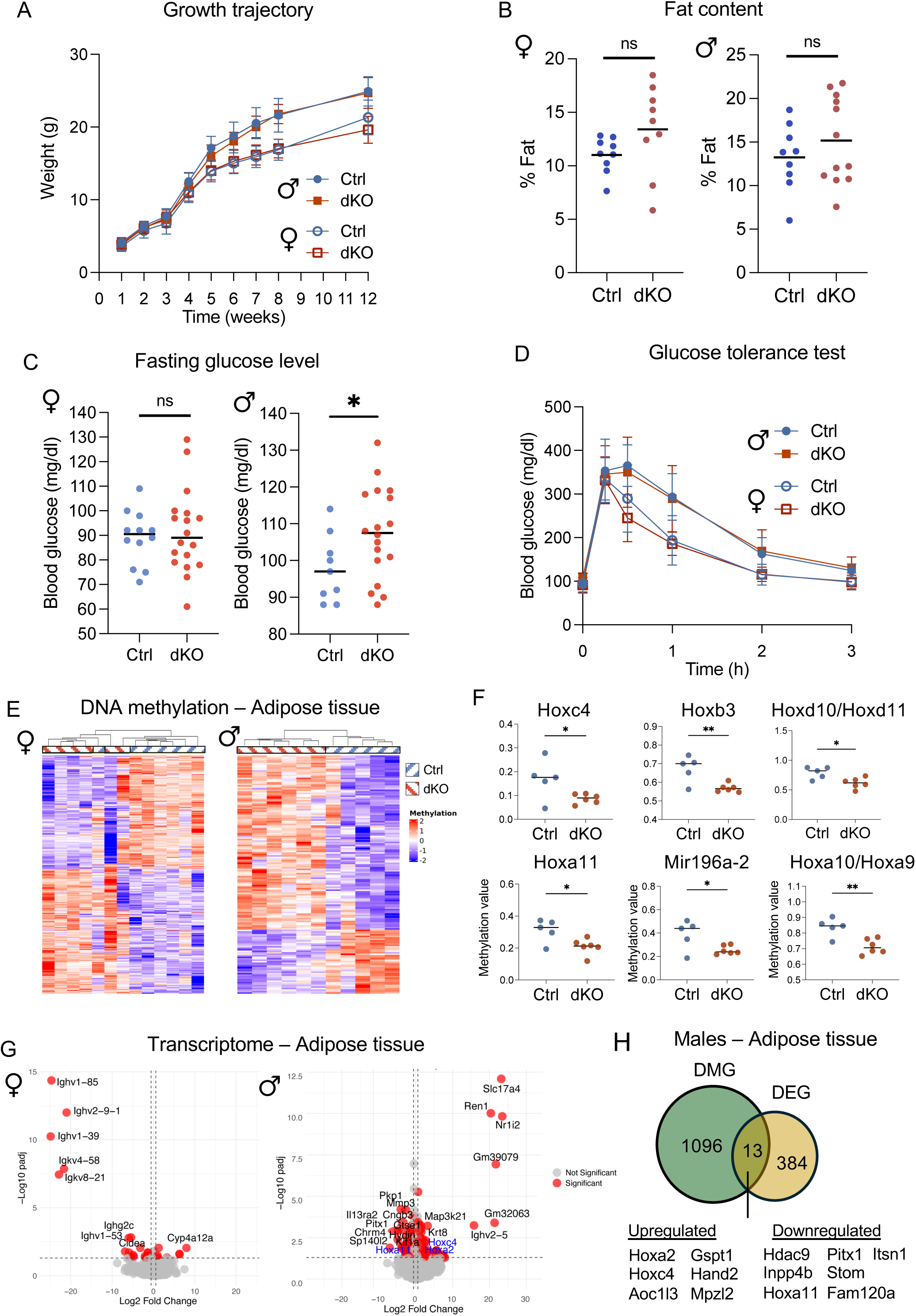
Increased Ca^2+^ at fertilization alters offspring metabolic parameters. (A) Growth trajectories of dKO derived pups and genotype matching controls (Ctrl) shown as mean weight ± SEM over time. N= 9 female and 9 male pups obtained from a minimum of 7 different control females (blue). N= 24 female and 30 male pups from 9 different dKO dams (red). (B-C) Percentage of fat content (B) and blood glucose levels after 14 h fast (C) in female (left) and male (right) pups derived from Ctrl (blue) or dKO (red) dams. Each dot represents an individual animal and horizontal bars indicate median. Mann Whitney test: *p<0.05 (D) Glucose tolerance test plotted as mean blood glucose ± SEM over time. N= 11 female and 8 male pups from a minimum of 7 different control females (blue). N=11 female and 14 male pups from 7 different dKO dams (red). (E) Heat map of differentially methylated probes found in adipose tissue of female (left) and male (right) offspring derived from Ctrl (blue stripes) or dKO (red stripes) dams. Methylation values are displayed as row-scaled β-values, ranging from −2 (indicating lower methylation, shown in blue) to +2 (indicating higher methylation, in red). (F) Dot plots of DNA methylation values (β-values) for six representative probes associated with Hox family members in adipose tissue from Ctrl (blue) and dKO (red) male-derived adipose samples. Unpaired t-test *p<0.05; **p<0.01. (G) Volcano plots of differential gene expression between dKO and Ctrl females (left) and males (right) in adipose tissue. Red dots indicate padj<0.05 while DEGs with padj<0.01 are labeled by name. Genes from the Hox family are highlighted in blue text. (H) Venn Diagram of DEGs and genes associated with differentially methylated probes (DMG) in adipose tissue from male dKO- and Ctrl-derived pups.

In various human and mouse studies, ART-derived offspring have abnormal DNA methylation in different tissues, affecting genes linked to adipocyte development^39,40^. Based on these studies, together with the abnormal glycemia and trend toward elevated fat content in the male dKO offspring, we evaluated the long-term impact of high Ca^2+^ exposure on DNA methylation in adipose tissue. Differentially methylated probes (DMP) were identified based on FDR<0.05 and average methylation difference >5% to capture relatively small but meaningful changes in methylation. In females, these parameters identified 513 DMP (244 hypomethylated and 269 hypermethylated probes), distributed across all genomic regions (Fig. 3E). Gene set enrichment analysis did not identify significant associations with specific biological functions. A similar analysis of male samples identified 1509 DMP (Fig. 3E). Of these, 520 were hypomethylated and 989 were hypermethylated in the dKO relative to controls. This high number of DMP led us to use a more stringent threshold of FDR<0.05 and methylation difference >10%, as it is generally accepted that a 10% change in DNA methylation can influence gene expression^41^. The new analysis identified 161 high-confidence DMP in males associated with 123 different genes, of which 36 were hypomethylated and 87 were hypermethylated. Molecular function analysis revealed a significant correlation between the 87 hypermethylation-associated genes and both Wnt signaling and tyrosine kinase activity, whereas the 36 hypomethylation-associated genes correlated with regulation of Pol-II transcription and DNA binding (Fig. S3C). Interestingly, 13 out of the 36 hypomethylation-associated genes belonged to the Homeobox (Hox) family, which is important in adipose tissue for fat distribution and function^42,43^. The Hox gene-associated probes were among the top candidates showing consistent differential methylation between dKO and Ctrl males, but not females (Fig. 3F).

Based on the DNA methylation results, we further investigated the impact of high Ca²⁺ exposure on RNA expression in the visceral adipose tissue of offspring. DEGs were defined by padj<0.05 and a fold change of ≥1.5. Similar to DNA methylation, females exhibited only few DEGs (29 DEG), whereas males had a significantly higher number, with 399 DEGs (Fig 3G). Functional analysis suggested that females had a deficiency in immune response (Fig S3D), while DEGs in males were associated with DNA binding, transcription, and retinoid X receptor (RXR) pathways (Fig S3E).

To determine if differential methylation in adipose tissue in males correlated with differential expression, an overlap of DEGs and genes associated with >5% DMP was performed. Thirteen genes overlapped, including three Hox genes and one histone deacetylase (Fig 3H). *Hoxa11*, *Hoxc4* and *Inpp4b*, all of which play a crucial role in metabolic regulation of adipose tissue, were included. These results indicate that differential methylation is likely to be a regulator of gene expression changes in Hox genes in male adipose tissue. Overall, these data demonstrate that high Ca^2+^ exposure shortly after fertilization has long term effects on male offspring metabolism and results in sexually dimorphic differential DNA methylation levels and gene transcription in visceral adipose tissue.

### 4. Excess Ca^2+^ influences H3K27ac and H3K18lac levels and global transcription in early embryos

Based on our results, we questioned how a brief metabolic insult at the 1C stage (lasting only hours) can impact embryo development and be amplified into a persistent metabolic phenotype in adulthood. To identify this link, we focused on changes occurring in the peri-fertilization window, when the epigenetic landscape is globally rearranged. Among the most abundant epigenetic modifications, H3K27ac and H3K18la are reorganized at the 1C and 2C stages and play key roles in EGA in both mice and humans^14,19^. In dKO embryos, H3K27ac levels were significantly increased at the 1C stage, about 8 h after fertilization, and this increase persisted at the 2C stage (Fig. 4A, 4B). In contrast, H3K18la levels in dKO embryos were significantly decreased in 1C embryos and remained so at the 2C stage (Fig. 4C, 4D). Global RNA synthesis as an indicator of EGA success was then compared between dKO and control embryos. dKO-derived embryos had extremely low global transcription at the mid-1C stage, which corresponds to the time of minor EGA (Fig. 4E). Transcription remained significantly reduced at the time of major EGA (2C stage; Fig. 4F), although the difference was less pronounced. These findings indicate that the dKO embryos had clear deficits in both minor and major EGA.

**Figure 4.**
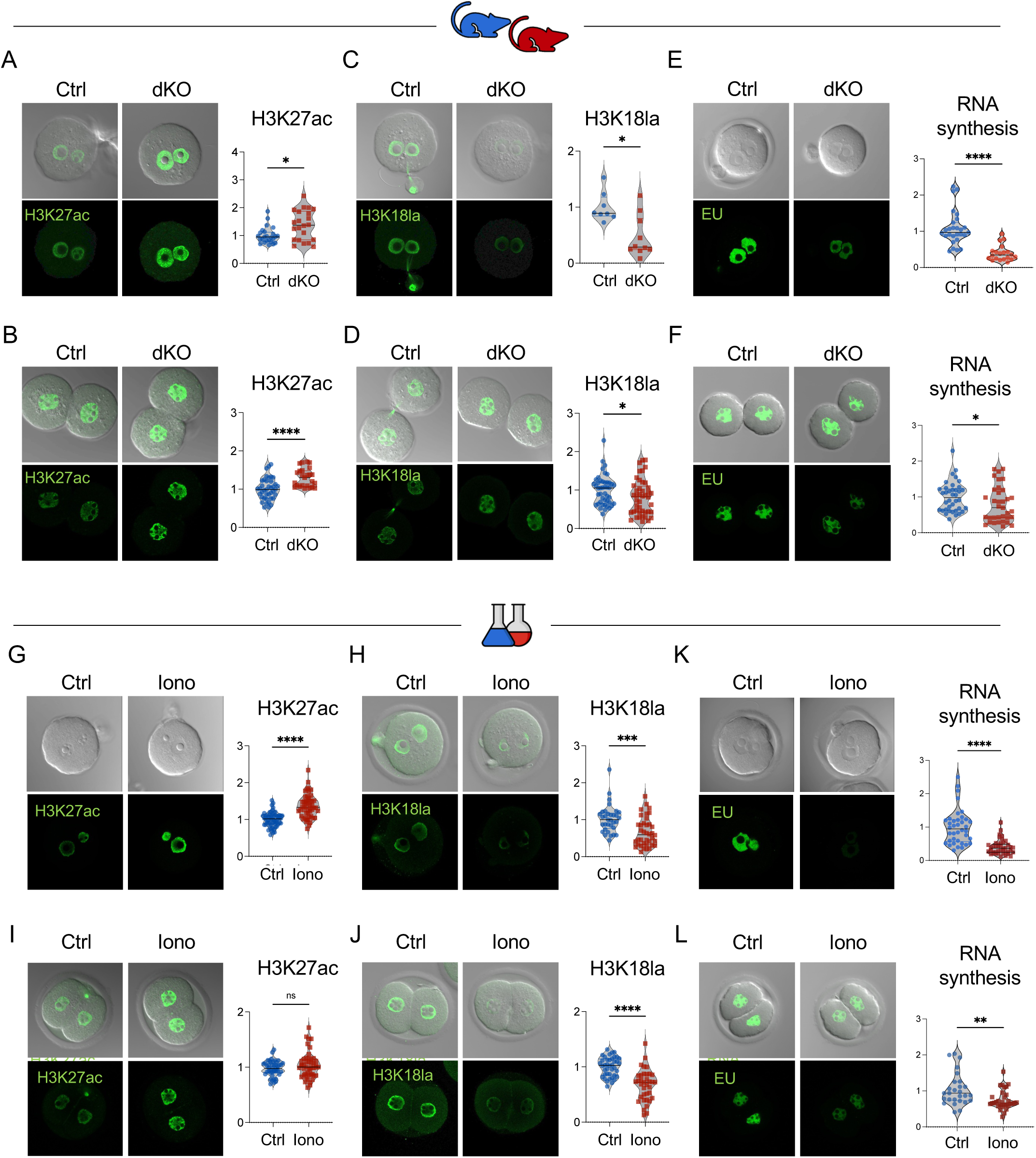
Excess Ca^2+^ influences H3K27ac and H3K18lac levels and global transcription in early embryos. (A-F) Representative images of nuclear staining of embryos obtained from Ctrl (blue) and dKO (red) mice, and corresponding quantification of the intensity relative to controls. H3K27ac levels at the 1C (A) and 2C (B) stages. H3K18la levels at the 1C (C) and 2C (D) stages. Global transcription as indicated by 5-ethynyl uridine (EU) staining at the 1C (E) and 2C (F) stages. (G-L) Representative images of nuclear staining and corresponding quantification of intensity relative to controls in wild-type embryos from control (Ctrl, blue) and ionomycin-treated (Iono, red) groups. (G) H3K27ac levels and (H) H3K18la levels at the 1C stage. (I) H3K27ac levels and (J) H3K18la levels at the 2C stage. EU staining at the 1C (K) and 2C (L) stages. Unpaired t-test was performed for datasets with a normal distribution (G, J). For all others, a Mann-Whitney test was used; *p<0.05, **p<0.01, ****p<0.0001; ns, not significant. Red and blue mouse icons represent different mouse models, while red and blue flasks indicate orthologous design using ionomycin.

One explanation for the EGA deficit was that the absence of PMCAs during oocyte growth and maturation somehow led to poor-quality oocytes, rather than just abnormal Ca^2+^ exposure at fertilization. To test this possibility, we performed an orthogonal assay where WT eggs were exposed to excess Ca^2+^ following fertilization using ionomycin; this treatment was done using an artificial oocyte activation protocol similar to those currently used in human ART clinics^12^. Like dKO embryos, ionomycin-treated embryos had increased H3K27ac levels and decreased H3K18la levels at the 1C stage relative to WT controls (Fig. 4G, 4H). By the 2C stage, H3K27ac levels returned to control levels, but H3K18la levels remained decreased, as observed in the dKO model (Fig. 4I, 4J). Excess Ca^2+^ exposure using ionomycin resulted in dramatically less global transcription at the 1C and 2C stages, again similar to the dKO derived embryos (Fig. 4K, 4L). The persistently low global transcription at the 2C stage in ionomycin-treated eggs, despite the restoration of H3K27ac levels to normal, suggests that H3K18la may play a more critical role regulating transcription. Altogether, these results demonstrate using two distinct methodologies, that excess Ca^2+^ exposure at fertilization regulates histone posttranslational modifications, leading to persistent changes at the 2C stage, and impairs minor and major EGA. Moreover, these findings show that ionomycin-treated WT eggs are a suitable model for abnormal Ca^2+^ exposure at fertilization, providing sufficient material for experiments that require larger numbers of embryos.

### 5. Excess Ca^2+^ at fertilization alters H3K27ac and H3K18la occupancy at the 2C stage

To determine the genomic locations where excess Ca^2+^ signals at fertilization impact H3K18la and H3K27ac marks, CUT&Tag was performed in control and ionomycin treated embryos after they reached the late 2C stage. A total of 10455 H3K18la peaks and 5155 H3K27ac peaks were identified in control samples. Over 94% of the H3K18la and H3K27ac peaks were located in distal regions or introns (Fig. 5A, 5B). There were 4443 differential H3K18la peaks and 4124 differential H3K27ac peaks between treated and control embryos (Fig. 5C). Consistent with the observed changes in immunofluorescence intensity, ionomycin exposure led to a net gain of about 500 H3K27ac peaks and a net loss of about 1500 H3K18la peaks. Ionomycin exposure primarily induced changes in H3K18la and H3K27ac peaks located in distal regions and introns (Fig. 5D, right), an unsurprising finding given the overall peak distribution. For H3K18la, a lower percentage of the peaks were located within 3 Kb of a transcription start site (TSS) compared to H3K27ac (Fig. 5D). Interestingly, the genomic location of differential H3K18la peaks largely did not correlate with that of differential H3K27ac peaks (Fig. 5C). The distinct locations and different responses to ionomycin suggest that Ca^2+^ triggered changes in H3K27ac and H3K18la are not mediated by the same chromatin modifiers.

**Figure 5.**
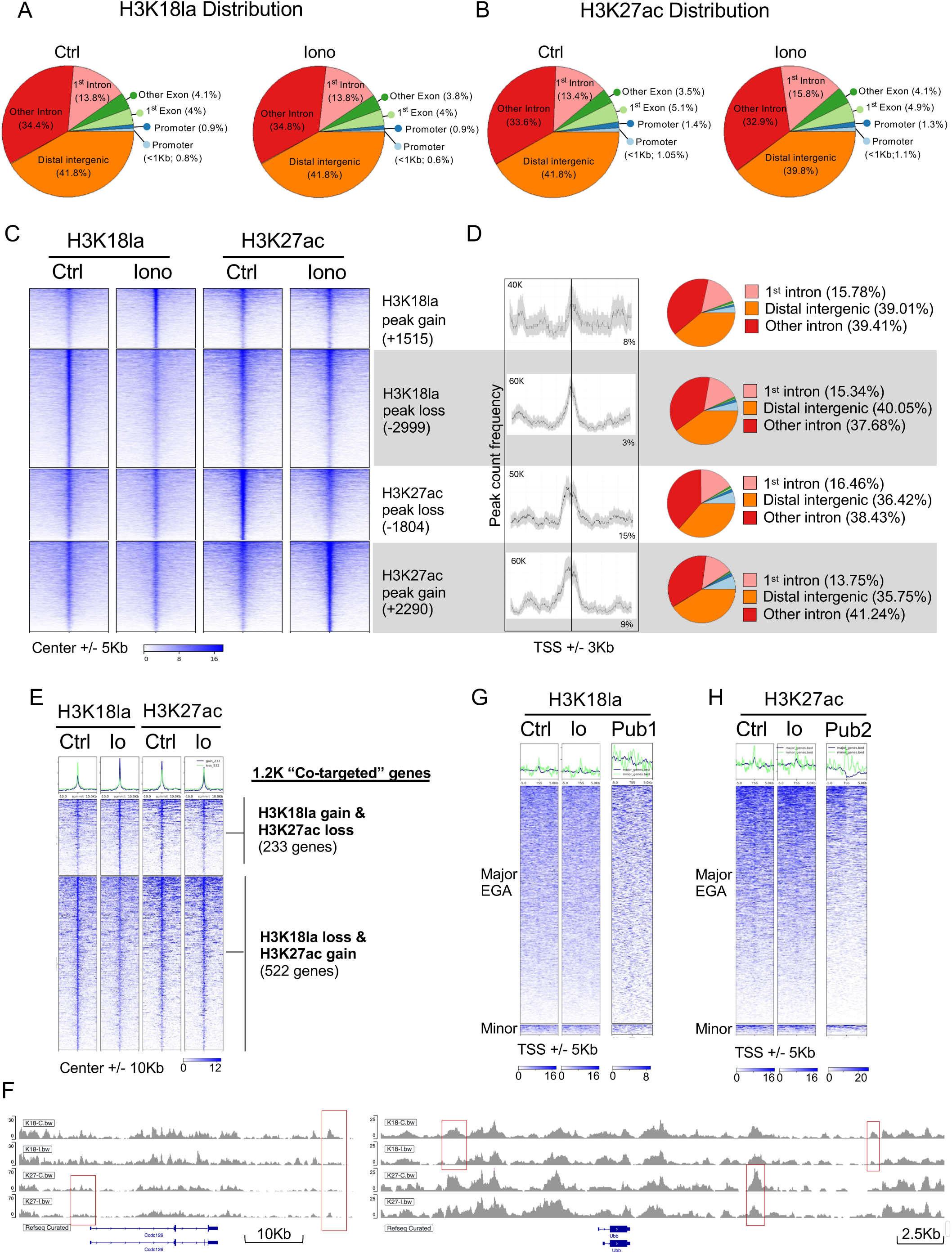
Excess Ca^2+^ at fertilization alters H3K27ac and H3K18la occupancy at the 2C stage. (A-B) Pie charts illustrating the genomic distribution of H3K18la (A) and H3K27ac (B) CUT&Tag marks in 2C-stage embryos. For each mark, distributions are shown for control (left) and ionomycin-treated embryos (right). (C) Heatmaps showing H3K18la and H3K27ac CUT&Tag signal in a 5 kb window around the summit, in control and ionomycin-treated embryos, organized by the highest to lowest signal. The number of gained and lost peaks after ionomycin treatment is indicated on the right. (D) Peak count frequency in promoters (+/− 3Kb from TSS) show minimal changes of H3K27ac and H3K18la marks in these regions. Pie charts on the right illustrate the genomic distribution of gained and lost peaks after ionomycin treatment. (E) Heatmaps and metaplots showing the co-occurrence of changes in H3K18la and H3K27ac marks within a close genomic region but in opposite directions, targeting the same associated genes (“co-targeted”). The top panel represents regions where H3K18la is gained while H3K27ac is lost, whereas the bottom panel shows regions where H3K18la is lost and H3K27ac is gained after ionomycin treatment. (F-G) Heatmaps and metaplots show changes in CUT&Tag signals for H3K18la (F) and H3K27ac (G) following ionomycin treatment at the promoters (TSS +/− 5Kb) of genes associated with major (top panel) and minor (bottom panel) EGA. Published reference data from Li et al. 2024^19^ (for H3K18la) and Li et al. 2022^18^ (for H3K27ac) are included for comparison. (H) Representative IGV tracks displaying CUT&Tag signals across two large genomic regions. Tracks show H3K18la signals in controls (top row) and ionomycin-treated samples (second row), as well as H3K27ac signals in controls (third row) and ionomycin-treated samples (fourth row). Differential peaks between ionomycin-treated and control samples are highlighted with red boxes.

To investigate the biological significance of Ca^2+^-mediated changes in H3K27ac and H3K18la occupancy, we identified the genes nearest to the differential peaks. There were 4364 genes linked to differential occupancy of H3K18la, and 2665 genes linked to differential occupancy of H3K27ac. Gene set enrichment analysis revealed no significant functional associations for either the gain or loss of H3K27ac or H3K18la peaks, suggesting that the observed changes in occupancy are not functionally coordinated. Among the genes associated with differential H3K27ac and H3K18la occupancy, 1238 genes had nearby regions occupied by both H3K18la and H3K27ac peaks; we refer to these as “co-targeted genes” (Fig. 5E, 5F). In 60% of the co-targeted genes, the occupancy of H3K27ac and H3K18la changed in opposite directions following ionomycin treatment: 233 genes gained H3K18la and lost H3K27ac occupancy, while 522 genes lost H3K18la and gained H3K27ac occupancy. No specific biological functions were enriched in these gene sets.

Because H3K27ac and H3K18la regulate mouse EGA, we examined whether EGA genes were associated with Ca^2+^-regulated differential peaks of either mark. In line with previous reports, a slight accumulation of H3K27ac and H3K18la was found at the promoters of major and minor EGA genes (Fig. 5G, 5H). However, ionomycin treatment did not cause any significant changes in H3K18la or H3K27ac occupancy at these genomic locations. Combined, these data revealed that excess Ca^2+^ significantly alters H3K27ac and H3K18la profiles in the early embryo, likely through independent mechanisms, but it does not impact their occupancy at promoters of EGA genes.

### 6. Excess Ca^2+^ at fertilization alters Pol-I-but not Pol-II-mediated transcription in early embryos

To better characterize the dramatic impact of Ca^2+^ on global transcription (Fig. 4K, 4L) we first estimated the relative contributions of Pol-I and Pol-II-mediated transcription to global transcription in our WT embryos. Inhibition of Pol-I alone by CX-5461 or inhibition of Pol-II alone by α-amanitin significantly reduced RNA synthesis at the 1C stage (Fig. 6A, 6B). Inhibition of both polymerases resulted in almost complete inhibition of RNA synthesis. There was no significant difference in the levels of Pol-I and Pol-II contributions to overall transcription, suggesting each contributes about 50% of the signal. Additionally, to assess whether there was a Ca^2+^ dose-dependent impact on transcription, two different conditions of high Ca^2+^ exposure were included: 1 pulse of ionomycin (Io1X), or our standard 2 pulses of ionomycin separated by 30 minutes (Io2X). A single pulse of ionomycin significantly reduced transcription to a similar extent as two pulses (Fig. 6A, 6B), indicating that even a slight increase in Ca^2+^ levels at fertilization was sufficient to reduce global transcription.

**Figure 6.**
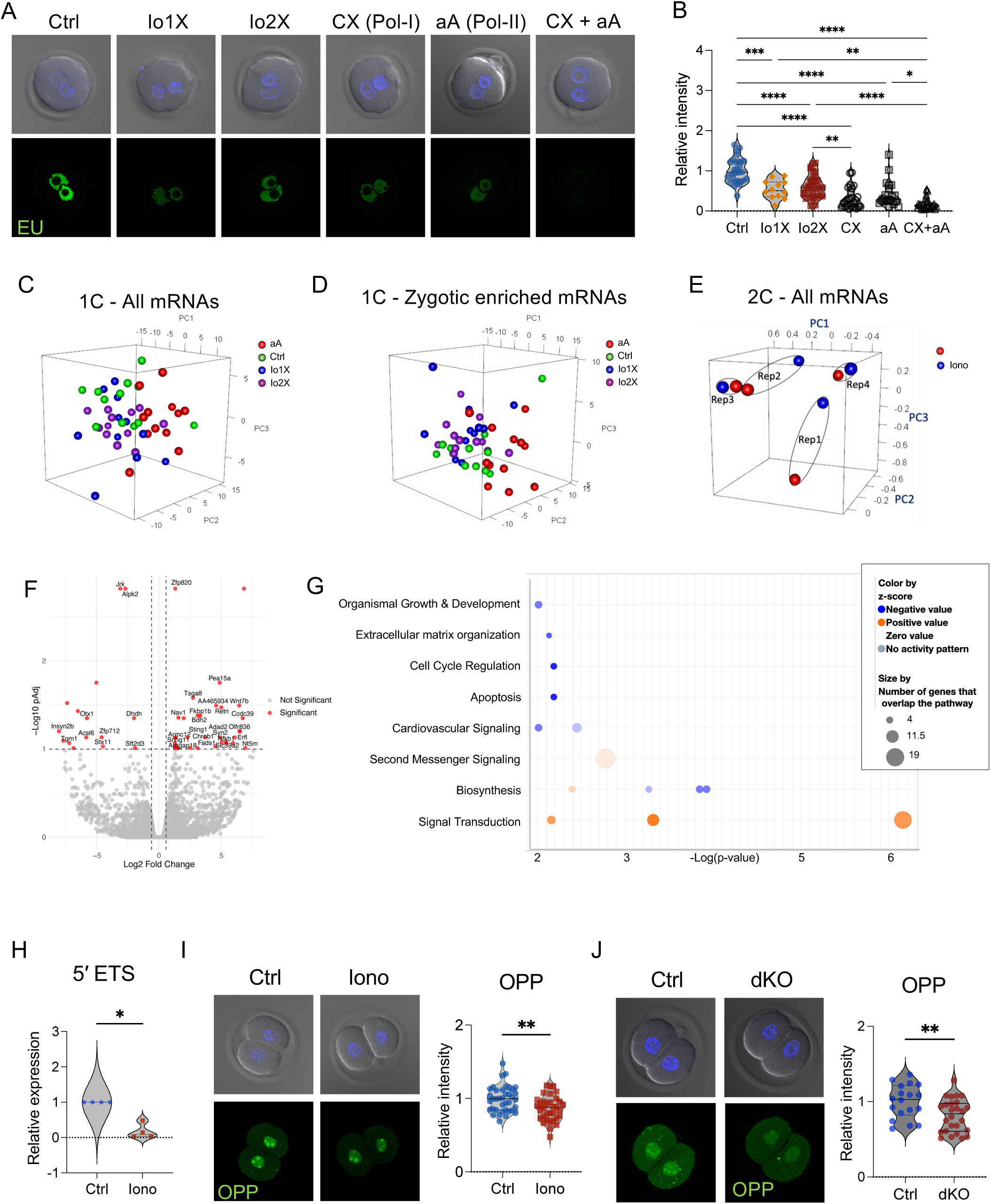
Excess Ca^2+^ at fertilization alters Pol-I-but not Pol-II-mediated transcription in early embryos. (A) Representative images of global transcription (EU staining) in 1C embryos and (B) corresponding quantification of nuclear staining. Ctrl, control; Io1X, 1 pulse of ionomycin; Io2X, 2 pulses of ionomycin; CX, inhibitor of RNA polymerase I (Pol-I); α-amanitin (aA), inhibitor of RNA polymerase II (Pol-II); CX+aA, combination of inhibitors targeting Pol-I and Pol-II. (C-E) Principal component analysis (PCA) from poly-A enriched RNAseq samples. (C) PCA plot of expression of all mRNAs detected in 1C embryos. (D) PCA plot of zygotic enriched mRNAs. Each dot indicates a single embryo. (E) PCA plot of expression of all mRNAs detected in late 2-cell (L2C) embryos. Each dot indicates a single biological replicate containing a pool of 5 embryos of Ctrl (blue) or Ionomycin (red) samples. (F) Volcano plot of differential gene expression between Ctrl and Iono-derived embryos at the late 2C stage. Red dots indicate padj<0.1. (G) Top canonical pathways from Ingenuity Pathway Analysis for DEGs in embryos at the L2C stage obtained from controls and ionomycin treatments. Canonical pathways are displayed on the Y-axis and are sorted according to their p-value. (H) Relative quantification of 5′ETS expression levels in Ctrl and Iono samples at the 2C stage. Each dot represents an independent biological replicate with at least 20 embryos. RT-qPCR data was analyzed by the ΔΔCT method using *Actb* (β-actin) as a housekeeping gene. Mann-Whitney test; *p<0.05. (I-J) O-propargyl-puromycin (OPP) staining showing new translation in (I) Ctrl vs Ionomycin or (J) Ctrl vs dKO embryos at the 2C stage. Graphs on the right show the quantification of nuclear signal relative to controls. Mann-Whitney test; **p<0.01.

To test the impact of excess Ca^2+^ on Pol-II, we performed poly-A enriched RNA sequencing (RNA-seq) in late 1C (minor EGA) and late 2C (major EGA) embryos. Newly synthesized RNAs were distinguished from stored maternal mRNAs by treating one group of embryos with α-amanitin; this group provided a concurrent dataset of 1C stage maternal mRNAs. At the late 1C stage, maternal and zygotic mRNAs were minimally affected by abnormal Ca^2+^ exposure, with only 33 differentially expressed genes (DEGs) (Fig. 6C). PCA analysis of zygotic enriched genes revealed incomplete separation between the groups (Fig. 6D). Only 13 minor EGA mRNAs were differentially expressed between control and excess Ca^2+^ groups, most of which were downregulated (*Zpf445, Gm12463, Aurkc, Mgme1, Dctn4, B3gntl1, Slc10a6, Scel, Adtrp, Egfr, Osbpl9, Orc3, and Arg2*). These minimal changes did not reflect the significant decrease in transcription at the 1C stage observed by immunofluorescence after high Ca^2+^ exposure (Fig. 4K), suggesting that Pol-I rather than Pol-II was affected. The impact of Ca^2+^-induced changes at the 2C stage on expression of major EGA mRNAs was similarly slight. PCA analysis revealed that the samples primarily segregated based on biological replicate rather than treatment (Fig. 6E). Only 51 genes were differentially expressed between groups; the most significant biological functions of these DEGs were G protein-coupled signal transduction and retinoic acid receptor activation (Fig. 6F, 6G). Only 10 out of the 51 DEGs were major EGA genes (*Cited1, Fkbp1b, Gm4340, Myo3a, Pglyrp1, Sat2, Sec61a2, Sparc, Tex101,* and *Tsga8*). Differential H3K18la peaks after ionomycin treatment were located near only 4 of the 51 DEGs (*Fads1, Nav1, Nt5m* and *Tsga8*) and only 3 had nearby differential H3K27ac peaks (*Acsl6, Nfkb1* and *Zfp712*). None of the DEGs showed simultaneously differential peaks for H3K18la and H3K27ac, suggesting that these histone marks are not directly responsible for regulating their expression.

To better understand whether non-polyadenylated RNA variants were affected by Ca^2+^, we focused on total RNA synthesis, and particularly Pol-I mediated transcription and ribosomal biogenesis. To assess Pol-I-mediated transcription, we quantified the expression levels of 5′ external transcribed spacer (5′ ETS), a region of the large ribosomal RNA precursor that is cleaved to form the mature rRNAs for ribosomal biosynthesis^44^. 5′ ETS levels were significantly reduced in ionomycin-treated 2C embryos (Fig. 6H), in line with the dramatically lower global transcription observed by immunofluorescence (Fig. 4K, 4L). Based on this result, we tested whether translation of RNAs into proteins was impacted by Ca^2+^. Indeed, global protein synthesis was decreased in both models of excess Ca^2+^ exposure, dKO-derived and ionomycin-treated 2C embryos (Fig. 6I, 6J). These findings strongly suggest that excess Ca^2+^ at fertilization impairs Pol-I mediated transcription and ribosomal biogenesis in early embryos.

### 7. Excess Ca^2+^ alters H3K27ac and H3K18la marks and global transcription by changing pyruvate flux

We next investigated the mechanism by which excess Ca^2+^ leads to alterations in H3K27ac and H3K18la, as well as reduced Pol-I-mediated transcription at the 1C and 2C stages. In early embryos, H3K27ac marks are primarily written by CREB-binding protein (CBP) and then erased by SIRT1^18,45–47^. A potential explanation for the alterations in H3K27ac was that excess Ca^2+^ affected nuclear localization of one or both of these proteins, but that was not the case (Fig. S4A, S4B). Because SIRT1 is a NAD+-dependent enzyme, its activity can be altered by changing the NAD+/NADH ratio. Based on the elevated levels of NADH in eggs exposed to high Ca^2+^ (Fig. 1E), NAD+ levels were likely to be reduced. To evaluate whether reduced availability of NAD+ following Ca^2+^ overexposure was responsible for differences in global transcription, control and ionomycin treated eggs were cultured with nicotinamide mononucleotide (NMN) to provide an external source of NAD+. Controls with or without NMN supplementation had comparable RNA synthesis at the 2C stage (Fig. S4C). As expected, zygotes treated with ionomycin had decreased global transcription compared to controls. However, ionomycin treated embryos had comparable levels of transcription regardless of whether they were supplemented with NMN, indicating that exogenous NAD+ was not sufficient to rescue the Ca^2+^-induced repression in RNA synthesis.

An alternative explanation was that Ca^2+^ alters the balance of acetyl-CoA and lactyl-CoA levels, which are essential for histone acetylation and lactylation, respectively. PDH catalyzes the irreversible conversion of pyruvate to acetyl-CoA and is one of the mitochondrial proteins translocated to the nucleus during EGA and cell reprogramming^32,33^. Interestingly, PDH activity is regulated by Ca^2+^ levels through its dephosphorylation, which is mediated by Ca^2+^-controlled pyruvate dehydrogenase phosphatase. Active PDH localized to the nucleus and was increased in ionomycin treated 2C embryos compared to controls (Fig. 7A), suggesting that excess Ca^2+^ promoted an increased flux from pyruvate to acetyl-CoA in this group. In line with this result, intracellular lactate levels were reduced in ionomycin-treated embryos, with levels about one-third of those detected in controls (Fig. 7B).

**Figure 7.**
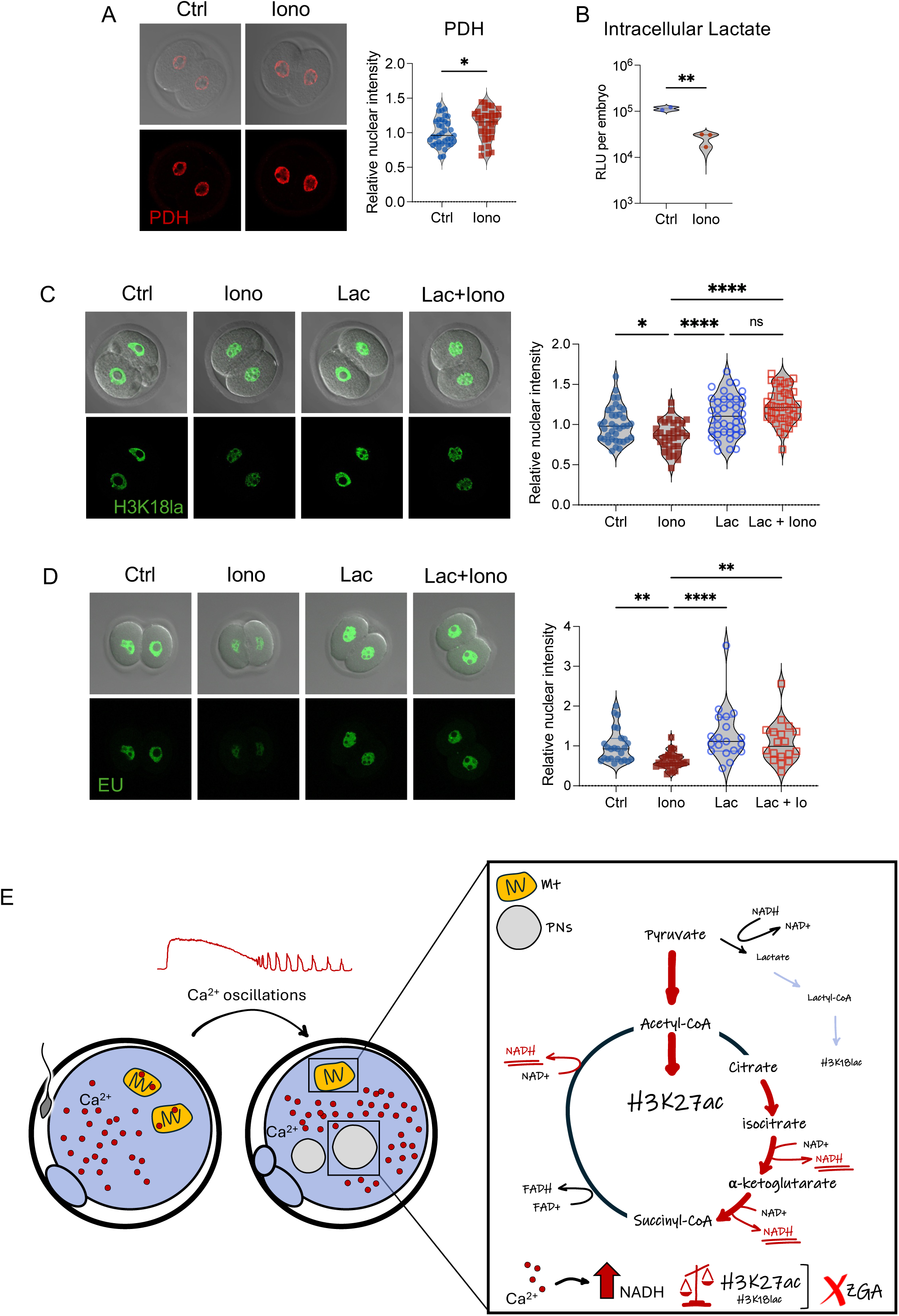
Excess Ca^2+^ alters H3K27ac and H3K18la marks and global transcription by changing pyruvate flux. (A) Representative images and quantification of nuclear staining for active pyruvate dehydrogenase (PDH) in control (Ctrl) and ionomycin-treated (Iono) embryos at the 2C stage. Unpaired t-test; *p<0.05. (B) Intracellular lactate levels at the 1C stage, expressed as relative light units (RLU). Mean is represented with a horizontal bar. T-test, **p<0.005. (C) Representative images and quantification of H3K18la levels in embryos at the 2C stage from Ctrl and Iono groups, as well as embryos microinjected with lactyl-CoA (Lac) and lactyl-CoA plus ionomycin (Lac+Iono). One-way ANOVA with Tukey’s multiple comparisons test; *p<0.05, ****p<0.0001; ns, not significant. (D) Representative images and quantification of global transcription activity (EU levels) in embryos at the 2C stage from Ctrl, Iono, Lac and Lac+Iono groups. Kruskal-Wallis with Dunnett’s multiple comparisons test; **p<0.005; ****p<0.0001. (E) Schematic representation of a working model illustrating Ca^2+^-triggered metabolic regulation of epigenetic reprogramming at fertilization.

We hypothesized that the increased flux from pyruvate to acetyl-CoA reduces the nuclear availability of pyruvate for conversion into lactate, leading to decreased lactyl-CoA levels and therefore reduced H3K18la. To test this idea, 1C embryos were microinjected with lactyl-CoA and then half were treated with ionomycin to increase the intracellular Ca^2+^ levels. Exogenous lactyl-CoA rescued the reduction in H3K18la levels caused by ionomycin treatment at the 1C and 2C stages (Fig. 7C, S4D). Although there was no impact on global RNA synthesis in controls, ionomycin treated embryos microinjected with lactyl-CoA had levels of RNA synthesis comparable to controls and significantly higher than uninjected ionomycin treated eggs (Fig. 7D). These results demonstrate that Ca^2+^-mediated alterations in H3K18la marks and RNA synthesis can be rescued by lactyl-CoA, but not NMN, and strongly suggest that restored levels of transcription after lactyl-CoA microinjection are due to restored H3K18la levels in the ionomycin treated group.

To further evaluate if excess Ca^2+^ exposure was associated with lower lactate levels (and therefore less lactyl-CoA), we compared our H3K18la differential peak locations in embryos exposed to excess Ca^2+^ to those from a published H3K18la CUT&Tag dataset generated from mouse 2C embryos deprived of lactate^19^. Interestingly, 24% of genes associated with differential H3K18la peaks overlapped, suggesting a shared pathway between lack of lactate and high Ca^2+^ exposure (Fig S4E). Altogether these findings suggest that the mechanism by which excess Ca^2+^ impairs placement of H3K18la marks is through changes in the endogenous availability of lactyl-CoA and subsequently, low H3K18la impairs global transcription in the early embryo.

## Discussion

In this study, we uncover one mechanism linking Ca^2+^ homeostasis at fertilization with developmental success, providing evidence for how zygote metabolism shapes offspring outcomes. We find that excess Ca^2+^ at fertilization impacts the embryo’s metabolic status and alters its epigenetic landscape and Pol-I-mediated transcriptional activity, thereby reducing total RNA synthesis—a phenomenon that persists at least until the 2C stage. Furthermore, we demonstrate that lactyl-CoA, but not an exogenous source of NAD+, is sufficient to mitigate the adverse effects observed at the 2C stage caused by excess Ca^2+^ at fertilization. Our findings strongly suggest that metabolic disruption is the primary driver of these outcomes, and lead to a model in which excess Ca^2+^ boosts the activity of mitochondrial enzymes, particularly PDH, which increases the availability of acetyl-CoA as a substrate for H3K27ac. The increased conversion of pyruvate to acetyl-CoA reduces its conversion to lactate, resulting in less lactyl-CoA for use a substrate for H3K18la, which impairs EGA (Fig 7E).

Previously, using a mouse model lacking only PMCA1 in the eggs, we showed that a 3-fold increase in Ca^2+^ at fertilization did not affect embryo development but had sexually dimorphic effects on offspring growth^5^. Here we show that a 10-fold increase in Ca^2+^ at fertilization impairs female fertility through its impact on preimplantation development and has sexually dimorphic effects on offspring metabolism. The more dramatic impact associated with the higher Ca^2+^ exposure supports a Ca^2+^ dose-dependent phenotype. However, using an in vitro approach, we found that even brief exposures to ionomycin after fertilization were sufficient to alter epigenetic marks and reduce global transcription, suggesting that even a slight increase in Ca^2+^ during this time-window is detrimental for embryo reprogramming. While our findings do not establish the full causal pathway linking Ca^2+^ changes at fertilization and long-term effects in the offspring, they suggest that Ca^2+^-induced epigenetic changes prime embryos for adult metabolic phenotypes. Our findings align with the previous demonstration that both a large excess and a large reduction in Ca^2+^ exposure at fertilization, produced by in vitro embryo manipulation followed by embryo transfer, differentially impact offspring growth and health^6,7,48^.

dKO derived males showed abnormal DNA methylation and gene expression in adipose tissue. Among the highly differentially methylated and expressed genes were *Hoxc4* and *Inpp4b*, both of which are involved in adipose function and metabolic regulation^43,49^. Abnormal *Hoxc4* expression in fat is sufficient to modulate metabolic function in mice and humans, while *Inpp4b*-KO males (but not females) have marked metabolic syndrome with higher adipose levels, insulin resistance, and inflammation in the adipose tissue^43,49^. Moreover, polymorphisms in mouse *HOXC4* are associated with abnormal distribution of body fat and other metabolic traits^43^. The long-lasting impact of altered Ca^2+^ exposure at fertilization on offspring adipose tissue provides one mechanism for the phenomenon of developmental origins of health and disease arising prior to implantation^50,51^.

To gain insights into the mechanism of Ca^2+^-mediated reprograming, we focused on metabolic changes in the peri-fertilization period. In line with Ca^2+^ enhancing TCA cycle activity, NADH levels were abnormally elevated in dKO eggs, changing the embryo redox status. In mice, adult weight is programed by the redox status (NADH/NAD^+^) as early as the 1C stage^7^. Also, in vitro produced embryos cultured in poorer quality conditions (media and oxygen tension) have lower levels of NAD+ at the blastocyst stage that correlates with lower developmental competence^52,53^. The key role of NAD+ at a very early stage of embryo development was demonstrated using an inhibitor of nicotinamide phosphoribosyltransferase, a key enzyme in NAD+ biosynthesis from nicotinamide^18^. Reduction of NAD+ from the 2C to 4C stage, but not from the 1C to 2C stage, resulted in failure to develop to the blastocyst stage, suggesting a strict NAD+ requirement at the 2C, but not the 1C stage. However, in this study the 1C embryos were flushed from the oviduct at a time in development after the Ca^2+^ oscillations cease and pronuclei form. These 1C embryos exhibited the highest levels of NAD+ among all preimplantation stages. Because NAD+ is a very dynamic metabolite that gets synthesized when needed for consumption (rather than accumulation), this finding suggests a significant requirement for NAD+ during the zygotic stage, potentially to support the high-energy demands associated with Ca^2+^ signaling during fertilization as well as epigenetic reprogramming. In line with this idea, the lack of NAD+ resulted in about 33% of minor EGA genes inappropriately upregulated at the 2C stage, while 25% of those persisted at the 4C stage, a clear impairment in minor EGA regulation^18^.

We found that Ca^2+^ induces changes in H3K27ac and H3K18la landscapes, as well as RNA synthesis, at the 1C stage, persisting to the 2C stage. Similarly, in immune cells, Ca^2+^ influx induces large-scale changes in H3K27ac and overall nuclear architecture, though H3K18la was not examined in that study^54^. Notably, our work shows that H3K18la and H3K27ac changes occur at different genomic locations, suggesting that they are mediated by different chromatin remodeling enzymes or scaffold proteins. During minor EGA, chromatin adopts a generally open state to permit broad, non-specific gene transcription. The widespread nature of H3K18la and H3K27ac changes without clear connections to functional pathways may simply reflect this open chromatin state. Interestingly, we found that the Ca^2+^-mediated effects on RNA synthesis were primarily linked to Pol-I. The requirement for Pol-I activity to support minor EGA and nucleolar formation was previously demonstrated in mice using a potent Pol-I inhibitor^22^. Therefore, the lesser effect of excess Ca^2+^ in dKO derived embryos on non-polyA RNA synthesis may explain why they have a drop in developmental potential after the 2C stage rather than a complete developmental arrest. Based on the previous results highlighting the importance of Pol I-related genes in minor EGA^22^, combined with our findings, we propose expanding the definition of minor EGA genes to include rRNAs.

To further confirm that disruptions in metabolism are the primary driver of Ca^2+^ mediated outcomes, we attempted to rescue phenotypic differences in embryos using different exogenous metabolites. On the one hand, Ca^2+^-induced abnormal reprograming, such as increased H3K27ac levels, could be attributed to impaired removal of this mark by SIRT1, due to low availability of NAD+. Supplying exogenous NAD+ was insufficient to rescue the reduced RNA synthesis, indicating that the lack of NAD+ was not mediating the observed effects. On the other hand, the Ca^2+^-induced increase in H3K27ac levels and decrease H3K18la levels could be explained by Ca^2+^-mediated allosteric activation of PDH. Increased PDH activity will enhance the irreversible conversion of pyruvate to acetyl-CoA as a substrate for H3K27ac and potentially reduce the conversion of pyruvate to lactate and lactyl-CoA as a substrate for H3K18la. Recently, in an elegant set of experiments it was demonstrated that lack of lactate negatively impacts H3K18la levels and leads to embryo arrest at the 2-cell stage^19^. Importantly, exogenous lactyl-CoA was sufficient to rescue this phenotype. In our hands, provision of exogenous lactyl-CoA before ionomycin treatment restored H3K18la levels at the 1C and 2C stages and successfully restored global transcription to control levels at the 2C stage. Moreover, there was a significant overlap between genes associated with differential peaks with/without lactate and with/without ionomycin, suggesting that ionomycin triggers a mild reduction in lactate availability within the nucleus. Of note, while the absence of lactate causes developmental arrest at the 2C stage, similar to the dKO embryo phenotype, the use of a low concentration of ionomycin does not impair embryo development, so an extensive overlap in between these datasets was not expected^55^.

Embryo requirements for metabolites dramatically change during preimplantation embryo development, with pyruvate and lactate required during the early stages and glucose after the 8C stage^51,56^. Embryo culture media, therefore, routinely include L-D lactate, raising the question of why L-D lactate in the culture media is not sufficient to mitigate Ca^2+^ mediated negative outcomes, while exogenous lactyl-CoA successfully does so. Our results strongly suggest that compartmentalized intracellular LDH activity (and potentially nuclear LDH) is crucial for embryo development and particularly for EGA. This agrees with previous reports in which 60% of mouse embryos cultured in medium containing both lactate and LDH inhibitors arrest at the 2C - 4C stage, the time of EGA^27^. Interestingly, although L-D lactate is supplied in the media throughout preimplantation development, it becomes highly enriched in the nuclei of mouse and human embryos during major EGA^19^. In mouse eggs, lactate-derived pyruvate does not enter the TCA cycle and instead may be metabolized through alternative pathways, one candidate being conversion of pyruvate to alanine^27^. This suggestion is based on findings that pyruvate contributes largely to energy production, but lactate-derived carbons are accumulated mainly in the protein fraction of 2C embryos^57^. Based on these and our results we propose that endogenous lactate produced by nuclear LDH is accumulated in the nucleus during EGA to support lactylation. Moreover, cytosolic lactate and lactate-derived pyruvate do not enter the TCA cycle but instead accumulate in the protein fraction, in the form of acetyl- or lactyl-marks, serving as a metabolic switch to drive changes for epigenetic reprogramming.

Newer highly sensitive techniques have revealed the critical roles of finely tuned and precisely timed metabolic changes in egg activation, preimplantation embryo development, and the developmental origin of health and disease, creating a new field of “Metabolofertility”. Our work advances this field by providing a mechanistic, metabolic connection between abnormal Ca^2+^ exposure, preimplantation embryo development, and offspring health. This connection is particularly relevant for ART clinics and especially for the indiscriminate use of ionomycin in the peri-fertilization window. Identifying this mechanism of Ca^2+^-mediated reprogramming is crucial because it creates opportunities to prevent and/or mitigate the negative outcomes associated with abnormal Ca^2+^ exposure at fertilization and, in the long term, will guide clinical therapies and protocols to improve offspring wellbeing.

## Author Contributions

V.S. and C.J.W. conceived the project and wrote the manuscript with input from all authors. V.S., P.S., and L.R. performed experiments and analyzed results. V.S. and M.A.E. designed experiments and analyzed results. D.D., B.N.P., Z.X. performed data analysis. C.J.W. supervised the project and secured funding.

## Acknowledgments

We kindly thank Dr. Jurien Dean for providing the hybridoma cell line (clone M2c.2) to obtain supernatant containing the anti-ZP2 antibody used in this study. We also thank Bob Petrovich and members of the Protein Expression Facility at NIEHS for production and purification of this antibody. This work was supported by the Intramural Research Program of the NIH, National Institutes of Environmental Health Sciences (1ZIAES102405) and by the NIH Pathway to Independence Award (1K99HD112510) supported by Eunice Kennedy Shriver National Institute of Child Health and Human Development (NICHD). The authors would like to thank the following NIEHS Cores for their support: Comparative Medicine Branch Animal Research Support (1ZIGES102585), Epigenomics and DNA Sequencing Core Facility (ZICES102545), Fluorescence Microscopy and Imaging Center (ZICES102485), Gene Editing and Mouse Model Core (ZICES102425), Integrative Bioinformatics Support Group (ZICES103371), Molecular Genomics Core (1ZICES102546) and the Protein Expression Facility (1ZICES102487).

## Methods

### Mice

C57BL/6J mice (strain #000664), and B6SJLF1/J males (strain #100012) were obtained from The Jackson Laboratory (BarHarbor, ME, USA). CF1 females (Hsd:NSA(CF1)) were obtained from Envigo (Indianapolis, IN, USA). A mouse model of dramatically increased Ca^2+^ exposure at fertilization in vivo was generated in house by targeting the *Atp2b1* and *Atp2b3* loci, encoding PMCA1 and PMCA3, the two main plasma membrane Ca^2+^ pumps in mouse eggs. The triple-target CRISPR method was used to target the *Atp2b3* locus in *Atp2b1* flox/flox (f/f) embryos previously generated in house^5,58,59^. Three single guide RNAs (sgRNAs) targeting *Atp2b3* exons 3 to 5 and a 200 bp single-stranded oligodeoxynucleotide (ssODN) homology-directed repair template carrying a triple stop (Table 1) were microinjected together with the CAS9 protein in *Atp2b1-f/f* 1-cell (1C) embryos about 20 h after hCG administration. Microinjected embryos were transferred at the 1C stage to oviducts of surrogate dams. Resulting founders were crossed to *Atp2b1-f/f*; Zp3-cre mice to generate dKO females globally null for PMCA3 and with an oocyte-specific deletion of PMCA1 *(Atp2b1-f/f*; Zp3-cre; *Atp2b3*-/-), and sibling females globally null only for PMCA3 *(Atp2b1-f/f*; *Atp2b3*-/-). To confirm the reproducibility of the phenotype, two independent mouse sub-lines derived from independent homology-directed repair events were generated (Figure S1). All founders were backcrossed for 4 generations with mice of *Atp2b1-f/f*; Zp3-cre background. Mice were maintained under controlled temperature and humidity, with a 12 h light/dark cycle. All animal procedures complied with National Institutes of Health animal care guidelines under an approved Animal Care and Use Committee protocol (RDBL07-37).

### Female fertility assessment

To determine the impact of Ca^2+^ levels on female fertility, twelve dKO females (*Atp2b1-f/f*; Zp3-cre; *Atp2b3*-/-), seven 3KO females (*Atp2b1-f/f*; *Atp2b3*-/-), and nine control females (*Atp2b1-f/f*; *Atp2b3*+/+) were mated to wild type (WT) males for 6 months. Parameters compared among groups were the numbers of litters, time between litters and number of pups weaned.

### Offspring genotypes, growth trajectory and metabolic assessment

The growth trajectories of control and dKO derived offspring were assessed as described before^5^. dKO females (*Atp2b1-f/f*; Zp3-cre; *Atp2b3*-/-) were mated to WT males to generate experimental offspring exposed to high Ca^2+^ levels at fertilization (female offspring: *Atp2b1*-/+; *Atp2b3*-/+; male offspring: *Atp2b1*-/+; *Atp2b3*-/y). To obtain genotype-matched control offspring exposed to physiological Ca^2+^ levels at fertilization, *Atp2b1*+/+; *Atp2b3*-/+ females were mated to *Atp2b1*-/+; *Atp2b3*-/y males. Of note, only ¼ female and ¼ male offspring carried the correct genotype and were included in the study. Pups were paw-tattooed 3 days after birth and weighed weekly for 8 weeks. At 3 weeks-of-age, the pups were weaned and placed on NIH-31 Rodent Diet. At 6 weeks-of-age, the mice were switched to an RD Western High Fat diet (D12079Bi, Research Diets Inc, New Brunswick, NJ, USA). Throughout the study, animals had ad libitum access to food and water. Weights were measured again at 12 weeks-of-age. Experimental mice and genotype-matched controls were assessed for glucose tolerance and body composition at 4 and 5 months of age, respectively. First, a sterile solution of D-glucose (250 mg/ml; Mallinckrodt Baker, Germany) was prepared. Glucose tolerance test (GTT) was performed in the morning, after a 14 h overnight fast with ad libitum access to water. Mice were weighed to calculate the proper amount of D-glucose to be injected, to ensure 2 mg D-glucose/g body weight. A sharp blade was used to snip the tip of the tail while the animals were briefly restrained, and the basal blood glucose levels were determined using a Nova Max® Plus glucometer (Nova Biomedical, Waltham, MA USA). The calculated amount of sterile D-glucose solution was administered by intraperitoneal injection.

Mice were placed back in their cages and blood samples were collected for plasma glucose measurements at 15, 30, 45, 60, 90, 120, and 150 minutes after the injection, from the lifted tail of the mouse. After the last measurement, mice were returned to their original cages. The iNSiGHT VET Dual-Energy X-ray Absorptiometry system (iNSiGHT VET DXA, OsteoSys, Guro-Gu, Seoul, Korea) was used for assessing body composition. A daily quality control check was performed following the manufacturer’s guidelines. Animals were weighed, sedated with vaporized 2.5% isoflurane in medical air using a nose cone and placed on the scanning table. Sedation was maintained throughout the assessment with 1.5% isoflurane using a nose cone inside the equipment chamber. Bone density and body composition, including fat weight and lean weight, were recorded for each animal. Following the manufacturer’s recommendation, X-ray images were preprocessed to hide the skull areas to remove interference. For that, a region of exclusion containing the skull was defined using the software provided by the manufacturer. After the measurements, mice were returned to their cages and monitored until recovery.

### DNA methylation analysis

The same control and experimental animals used to test growth trajectories were assessed for differences in DNA methylation in adipose tissue. At 6 months of age, adipose tissue samples were collected, snap-frozen in dry ice and stored at −80°C until use. Genomic DNA was isolated from 50 mg of each tissue. Bisulfite conversion was performed using the EZ DNA Methylation-Lightning Automation kit (Zymo Research, Irvine, CA, USA) with 820 ng starting DNA. DNA analysis was conducted using Infinium Mouse Methylation BeadChip arrays (Illumina, San Diego, CA, USA) following the Illumina InfiniumHD methylation protocol. Starting with ~4 µl of bisulfite converted DNA, samples were amplified for 21 h at 37°C using random primers. The amplified DNA was then fragmented, precipitated, and resuspended prior to hybridization to BeadChips for 18 h at 48°C. The slides were washed, stained, and then scanned with an Illumina iScan.

The Infinium Mouse Methylation BeadChip data were preprocessed using the ENmix software, which included ENmix background correction^60^, RELIC dye bias correction^61^, quantile normalization of intensity values separately for type I and type II probes, and RCP probe type bias adjustment^62^. Outlier values for each probe, defined as those exceeding three times the interquartile range, were excluded, and missing values were imputed using the KNN algorithm^63^. CpG probes with more than 5% low-quality data were removed from the analysis. Low-quality data were defined as data points with fewer than three beads or a detection p-value greater than 10⁻⁶. After preprocessing and quality control, 72 samples and 279,522 probes remained. A robust linear regression model was used to assess the treatment effect on long-term DNA methylation changes at each individual CpG site, stratified by sex. A false discovery rate of 0.05 and an average methylation level difference of either 5% or 10% between treatment and control groups were used as significance thresholds. All analyses were conducted using R version 4.3.3.

### Gamete and embryo collection

Female mice (age 6-12 weeks) were superovulated as described before^64^. Briefly, for germinal vesicle (GV) stage oocyte collection, females were injected intraperitoneally with 7.5 IU of equine chorionic gonadotropin (eCG, Lee Biosolutions, Maryland Heights, MO, USA) and ovaries collected 44-48 h after injection. The ovaries were placed in pre-warmed Minimal Essential Medium (MEM) with Hepes (Thermo Fisher, Waltham, MA, USA) containing 0.1% polyvinyl alcohol (PVA; Millipore Sigma, St. Louis, MO, USA) and 2.5 μM milrinone (Millipore Sigma) to prevent resumption of meiosis. GV oocytes surrounded by at least 1 complete layer of cumulus cells were selected and placed in drops of MEM α (Thermo Fisher) supplemented with 5% FBS (10438018, Thermo Fisher) and 2.5 μM milrinone under mineral oil in a humidified atmosphere of 5% CO_2_ at 37°C until use. For collection of eggs (metaphase II-arrested eggs), females were injected with eCG, followed 44-48 h later with 7.5 IU of human chorionic gonadotropin (hCG; Millipore Sigma). At 13-16 h after hCG administration, oviducts were collected in pre-warmed MEM with Hepes containing 0.1% PVA. When necessary, cumulus cells were removed in MEM containing HEPES, 0.1% PVA and 0.1% hyaluronidase (Millipore Sigma). Denuded MII eggs were placed in drops of human tubal fluid medium (HTF; Millipore Sigma) containing 3 mg/ml of AlbuMax I lipid rich BSA (Thermo Fisher) (HTF-BSA) under mineral oil in a humidified atmosphere of 37°C and 5% CO_2_ until needed. For 1C embryo collection, females were superovulated as described before and mated with B6SJLF1/J males of proven fertility immediately after hCG injection. 1C embryos were collected 17-20 h after hCG administration, in the same way as MII eggs.

### In vitro fertilization (IVF), ionomycin treatment, and embryo culture

IVF was performed using fresh sperm as previously described^5^. Briefly, epididymides were collected from an adult B6SJLF1/J male and placed in a 500-µL drop of HTF-BSA medium under mineral oil. Using forceps, 4–5 snips were made to allow the sperm to swim out to the media. After 15 min, 150 µL of the sperm-containing medium was transferred to a tube containing 850 µL of HTF-BSA and incubated vertically to allow motile sperm to swim up. After 60 min, the middle layer of the medium, containing the motile sperm, was collected and the sperm were counted. The IVF dish was prepared with 300-µL drops of HTF-BSA containing 5×10^5^ sperm/mL, covered with mineral oil. Oviducts from superovulated CF1 female mice were collected, and the ampullae were opened with a needle to release the cumulus-oocyte complexes (COCs) directly into the sperm-containing drops. IVF was conducted for 3 hours. Presumptive zygotes were washed free of unbound sperm, unfertilized eggs removed, and the zygotes divided randomly into the different experimental groups. For ionomycin treatment, immediately after IVF, presumptive 1C embryos were placed in KSOM medium containing 2 µM ionomycin (I0634; Millipore Sigma) for 5 min, then washed in KSOM. The same ionomycin treatment was repeated after 30 min. Finally, embryos were cultured in KSOM medium covered with mineral oil at a ratio of 1 embryo/µL KSOM in a humidified atmosphere of 5% CO_2_ and 5% O_2_ at 37°C. For zona-free IVF, eggs were cultured for 3-6 s in drops of acid Tyrode’s solution (pH=1.6) to remove the zona pellucida and washed in 5 50-µl drops of KSOM. For fertilization, 10 zona-free eggs were placed in 10-µL drops of HTF-BSA containing 1×10^4^ sperm/mL, covered with mineral oil. Zona-free IVF was conducted for 1 h. Embryos were then washed and cultured singly in 1 µL drops of KSOM medium under mineral oil in a humidified atmosphere of 5% CO_2_ and 5% O_2_ at 37°C.

### Ratiometric Ca^2+^ imaging

Zona-free eggs were loaded with a Ca^2+^ dye by incubation in KSOM (Millipore Sigma) containing 0.02% Pluronic F-127 (Thermo Fisher) and 5 µM Fura-2LR/AM (Thermo Fisher) for 30 min. Control, 3KO, and dKO eggs were then placed side by side in a 150 µL drop of KSOM without BSA on a glass-bottom dish (MatTek, Ashland, MA) coated with Cell-Tak (Millipore Sigma). Once the eggs adhered to the dish, 45 µL of HTF medium supplemented with 3 mg/mL AlbuMax I lipid-rich BSA (Thermo Fisher) was added, and the medium was covered with mineral oil. Ratiometric Ca^2+^ imaging was performed as described previously^64^. Baseline fluorescence was recorded, and 3-5 min later, the dish was inseminated to achieve a final sperm concentration of 1×10^4^ sperm/mL. Ca^2+^ levels were recorded every 7.5 seconds using a Hamamatsu ORCA-Flash 4.0 LT+ digital camera (Hamamatsu, Bridgewater, NJ, USA) mounted on a Nikon Ti inverted microscope. The system was equipped with a Nikon S Fluor 20×/0.75 NA objective (Nikon Instruments, Melville, NY, USA), and excitation wavelengths of 340 nm and 380 nm were applied using a Lambda 10-B Optical Filter Changer (Sutter Instruments, Novato, CA, USA). Ca^2+^ levels were expressed as a ratio of fluorescence signal (F340/380). To evaluate total intracellular Ca^2+^ stores, Fura-2-loaded eggs were placed in 1800 µL of Ca^2+^- and magnesium-free CZB medium (CMF-CZB) without BSA. After recording baseline fluorescence for 3-5 minutes, 200 µL of ionomycin prepared in CMF-CZB was added to achieve a final concentration of 5 µM. Intracellular Ca^2+^ imaging was performed following the same procedure described for IVF. Relative internal Ca^2+^ stores between controls and dKO were assessed by comparing the peak amplitude immediately after ionomycin treatment. This approach avoided distortion of the Ca^2+^ store measurements due to impaired cytoplasmic Ca^2+^ efflux in dKO embryos, which lack the two major Ca^2+^ efflux pumps.

### Simultaneous imaging of Ca^2+^ and NADH levels at fertilization

Ca^2+^ imaging was performed as described before, but using Fura Red-AM (F3021, Thermo Fisher). For Ca^2+^ detection, the 458 nm and 488 nm lines from an Argon laser were used for excitation. Emission was collected through a 596-758nm band pass filter. For NADH detection, UV light (355 nm UV laser) was used for excitation, while emission was collected using a 435-486 nm long pass filter as previously described^30^. Signals were recorded every 15 sec using a Zeiss LSM 780 inverted confocal microscope equipped with a stage incubator with 5% CO_2_ and temperature controlled at 37°C.

### Inhibition of transcription

Immediately after IVF, fertilized eggs were cultured in 96-well plates with 60 µL KSOM medium containing specific inhibitors. Pol-II transcription was blocked using 24 µg/mL α-amanitin (Sigma-Aldrich, A2263)^65^. Pol-I was inhibited with 0.5 mM CX-5461^22^ (Millipore Sigma). Both inhibitors were combined when necessary. CX-5461 was used without an oil overlay to avoid concentration alterations^66^. Embryos were cultured at 37°C in a humidified atmosphere of 5% CO_2_ and 5% O_2_ at 37°C.

### Rescue experiments

Nicotinamide mononucleotide (NMN, N3501; Millipore Sigma) was prepared in autoclaved PicoPure water to a stock concentration of 1 mM. Lactyl-CoA (HY-141540, MedChemExpress, Junction, NJ, USA) was prepared to 50 mM in DMSO. Immediately after IVF, fertilized eggs were cultured in KSOM medium supplemented with 1 µM NMN as previously described^18^. After 30 min, a subset of NMN-treated eggs was exposed to ionomycin as described before and returned to KSOM containing NMN. Eight h after initiation of IVF, zygotes were washed and transferred to regular KSOM. Control groups included fertilized eggs with or without ionomycin treatment with no NMN supplementation. Lactyl-CoA was diluted in PBS to a final concentration of 0.2 µM. For microinjection experiments, IVF was performed for 1.5 h. A group of fertilized eggs was microinjected with about 10 pL of lactyl-CoA immediately after IVF using an inverted microscope equipped with TransferMan 4r micromanipulators (Eppendorf North America, Framingham, MA, USA) and an Eppendorf FemtoJet microinjector. Microinjected zygotes were cultured in KSOM. Three hours post-fertilization, a subset of lactyl-CoA-injected eggs was treated twice with ionomycin, following the same protocol. Uninjected embryos with and without ionomycin treatment served as controls.

### Immunofluorescence

Embryos were fixed with 2.5% paraformaldehyde in PBS for 30 min and then washed in PBS containing 1 mg/ml BSA and 0.01% Tween-20 (blocking solution). Permeabilization was performed for 15 min using 0.2% Triton X-100 in PBS. Immediately after permeabilization, embryos were incubated at room temperature (RT) for 1 h in blocking solution, followed by overnight incubation at 4°C with a 1:100 dilution of the primary antibody. After three washes in blocking solution (15 min each), embryos were incubated for 1 h at RT with a 1:500 dilution of a fluorochrome-conjugated secondary antibody. Because anti-γH2AX is a conjugated primary antibody, no secondary antibody incubation was performed. Finally, embryos were mounted on a slide with Vectashield containing 1.5 μg/mL DAPI (Vector Laboratories, Burlingame, CA, USA) and covered with a coverslip for imaging. Slides were scanned using a Zeiss LSM 780 inverted confocal microscope. The list of primary and secondary antibodies is in Table 1.

For quantification of nuclear staining, images were opened in FIJI/ImageJ, and a region of interest (ROI) was drawn around each pronucleus in 1C embryos or each nucleus in 2C embryos to measure the average pixel intensity. The values obtained for each ROI were averaged to determine the intensity per embryo. All values were normalized relative to the average of the corresponding controls.

### Cortical granule staining

Eggs were fixed with 4% paraformaldehyde in PBS for 60 min and then washed for 15 min in PBS containing 3 mg/ml BSA (PBS/BSA) and 0.1M glycine. They were then permeabilized in PBS/BSA containing 0.1% Triton X-100 for 15 min. Following a 15 min wash in PBS/BSA containing 0.01% Tween-20 (P/B/Tw), eggs were incubated for 30 min in P/B/Tw containing 10 µg/mL fluorescein-conjugated lens culinaris agglutinin (FL-1041-5; Vector Laboratories). The eggs were washed 3 times for 15 min each in P/B/Tw, and mounted on slides with DAPI-containing vectashield, as described above.

### Immunoblotting

For ZP2 detection, 10 GV oocytes, MII eggs, and 2C embryos were collected, snap-frozen on dry ice, and stored at −80°C. Upon thawing, 10 µL of sample buffer was added to each sample, and the mixture was heated for 5 min at 99°C before being placed on ice. Samples were loaded onto a 4–20% Bio-Rad gradient Tris-Glycine gel, and proteins were transferred to a PVDF membrane using the Trans-Blot® Transfer System™ (Bio-Rad, Hercules, CA, USA). Blots were blocked for 1 h at RT in blocking solution (5% non-fat dry milk in TBST) and incubated overnight at 4°C with a 1:5000 dilution of anti-ZP2 (clone M2c.2)^67^ supernatant in blocking solution. Blots were washed three times with TBST and incubated for 1 h at RT with a 1:10,000 dilution of peroxidase-conjugated anti-rat IgG, while gently agitating. For signal detection, a 1:4 dilution of SuperSignal™ West Femto Maximum Sensitivity Substrate (Thermo Scientific) was applied, and images were captured using the Bio-Rad ChemiDoc™ Touch Imaging System.

### Lactate assay

For intracellular lactate measurement, 8 h after IVF, pools of 50 to 74 1C embryos were washed three times in PBS containing PVP to remove extracellular lactate. Samples were then snap-frozen in a minimal volume and stored at −80°C until use. For each replicate, an equal number of control and ionomycin-treated embryos were collected. On the day of the assay, embryos were thawed on ice. The Lactate-Glo™ Assay (Promega, Madison, WI, USA) was performed following the manufacturer’s instructions for tissue. A premix was prepared using an 8:1 ratio of homogenization buffer (50 mM Tris, pH 7.5) to inactivation solution (0.6 N HCl). Each sample was treated with 50 µL of this premix, followed by the addition of 5 µL of inactivation solution (1 M Tris, pH 10.7). Next, 50 µL of each prepared sample was transferred to a microplate reader, and 50 µL of Lactate Detection Reagent was added. The plate was incubated at RT in the dark for 1 h before luminescence was recorded using a GloMax® Luminometer (Promega). Data are presented as the average relative light units (RLU) per embryo.

### CUT&Tag, library construction and sequencing

CUT&Tag for H3K27ac and H3K18la was performed using the Hyperactive Universal CUT&Tag Assay Kit for Illumina Pro (TD904, Vazyme, Nanjing, China), following the manufacturer’s instructions with slight modifications. Briefly, for H3K27ac, 42 2C embryos were lightly fixed in 0.1% paraformaldehyde for 1 min at RT. After 2 washes in wash buffer, the embryos were incubated with concanavalin A-coated magnetic beads for 10 min at RT. For H3K18la, no fixation was performed. Instead, 50 embryos were directly incubated with the concanavalin A-coated magnetic beads, as described above. The samples were then incubated with a 1:50 dilution of the primary antibody (rabbit polyclonal anti-H3K27ac, Active Motif, Carlsbad, CA, USA, cat # 39133, or rabbit monoclonal anti-H3K18la, PTM Bio, Chicago, IL, USA, cat # PTM-1427RM) overnight at 4°C. After washing twice with Dig-wash buffer, they were incubated in a 1:100 dilution of guinea pig anti-rabbit IgG (Antibodies Online, Limerick, PA, USA, cat # ABIN101961) for 1 h at RT, followed by transposase fused with protein A/G (pA/G-Tnp Pro) for 1 h at RT. Fragmentation and DNA extraction were performed following the manufacturer’s instructions, including the addition of 1 fg of the spike-in DNA provided in the kit. For library amplification, 20 cycles of PCR were done using the running parameters suggested in the kit, and including indexes from the TruePrep Index Kit V2 for Illumina (Vazyme, cat # TD202). Purification of PCR products was performed using VAHTS DNA Clean Beads (Vazyme, cat # N411) and pooled libraries were sequenced on the Illumina NextSeq platform to generate 50 bp paired-end reads.

### CUT&Tag analysis

Paired-end fastq files were aligned to the mouse mm10 reference genome with bowtie2 version 2.5.2 (Langmead and Salzberg, 2012) after removing low quality reads with fastp (Chen et al., 2018). The alignment parameters were (bowtie2 -p 8 -X 700 --local --very-sensitive-local --no-unal --no-mixed --no-discordant --phred33 -I 10). Aligned .sam files were converted to coordinate sorted binary .bam files with picard SortSam. Normalized bigwig read coverage files were created using deepTools bamCoverage version 3.5.1^68^. Peaks were called with MACS2 callpeak version 2.2.9.1^69^ using parameters (-p 1e-5 -f BAMPE -g hs --keep-dup all) for paired-end reads. Differential peaks were determined with MAnorm version 1.3.0^70^. Heatmaps were generated using deepTools.

### RNA isolation, cDNA synthesis and RT-qPCR

Pools of 20–50 2C embryos were collected, flash-frozen on dry ice, and stored at −80°C. Equal numbers of control and treated embryos were collected to minimize variability in RNA isolation efficiency. RNA was extracted using the Arcturus PicoPure RNA Isolation kit (Thermo Fisher) according to the manufacturer’s instructions and eluted in 11 µL of elution solution. Complementary DNA (cDNA) synthesis was performed immediately after RNA isolation using 10 µL of RNA as input and the SuperScript First-Strand Synthesis System for RT-PCR (Thermo Fisher), as previously described^71^. qPCR was performed using one embryo equivalent of cDNA. The ΔΔCt method was used to calculate the relative expression of 5′ETS using *Actb* as reference^72^.

### Poly-A enriched RNA library preparation

Single 1C embryos (14 h after IVF) or pools of 5 late 2C embryos (36 h after IVF) were flash-frozen on dry ice in minimal media and stored at −80°C. The Pol-II inhibitor α-amanitin (aA) was used to suppress transcription in a group of 1C embryos immediately after IVF. This group enable the identification of maternally inherited mRNAs and the discrimination of zygotically enriched mRNAs. cDNA synthesis was performed using the SMART-Seq® v4 Ultra® Low Input RNA Kit (Takara, San Jose, CA, USA, 634888) according to the manufacturer’s instructions, with 15 cycles of amplification. Amplified cDNA was cleaned using Agencourt AMPure XP beads (Beckman Coulter, Brea, CA, USA, A63881) and eluted in 17 µL. Libraries were prepared with the Nextera XT Kit (Nextera, Juno Beach, FL, USA) using 1 ng of input cDNA and the Nextera® XT Index Kit v2 (Set A). Libraries were cleaned using Illumina Purification Beads (IPB) and resuspended in 30 µL resuspension buffer. Library quality was assessed on an Agilent Bioanalyzer using a High Sensitivity DNA Kit, and quantification was performed using a Qubit 3.0 Fluorometer (Thermo Fisher). Pooled libraries were sequenced on an Illumina NovaSeq 6000 NGS system to generate 50 bp paired-end reads.

### RNAseq analysis

RNAseq fastq files were analyzed with fastp version 0.23.4 to remove adapters and low-quality reads^73^. The resulting files were then aligned to the mm10 mouse reference genome with STAR version 2.6.0c ^74^. On average, 34 million uniquely aligned reads were obtained per sample. Gene read counts and annotations were obtained with the R subread package, version 1.28.1^75^. To determine differentially expressed genes, we used the R package DESeq2 version 1.42.1^76^ to compare ionomycin and control samples. Heatmaps and 3D PCA plots were made using R packages, “pheatmap” and “rgl”, respectively. For comparison, previously reported maternal and ZGA gene lists were used^77^.

### Statistics

Except for DNA methylation and NGS data, all statistics were performed using GraphPad Prism, v10. Normality and lognormality tests were initially performed on each dataset. Accordingly, two-tailed t-tests or Mann Whitney tests were used for parametric or non-parametric data. Similarly, ANOVA or Kruskal-Wallis tests were performed for normally distributed or non-parametric data sets containing more than 2 groups.

## Key resources table

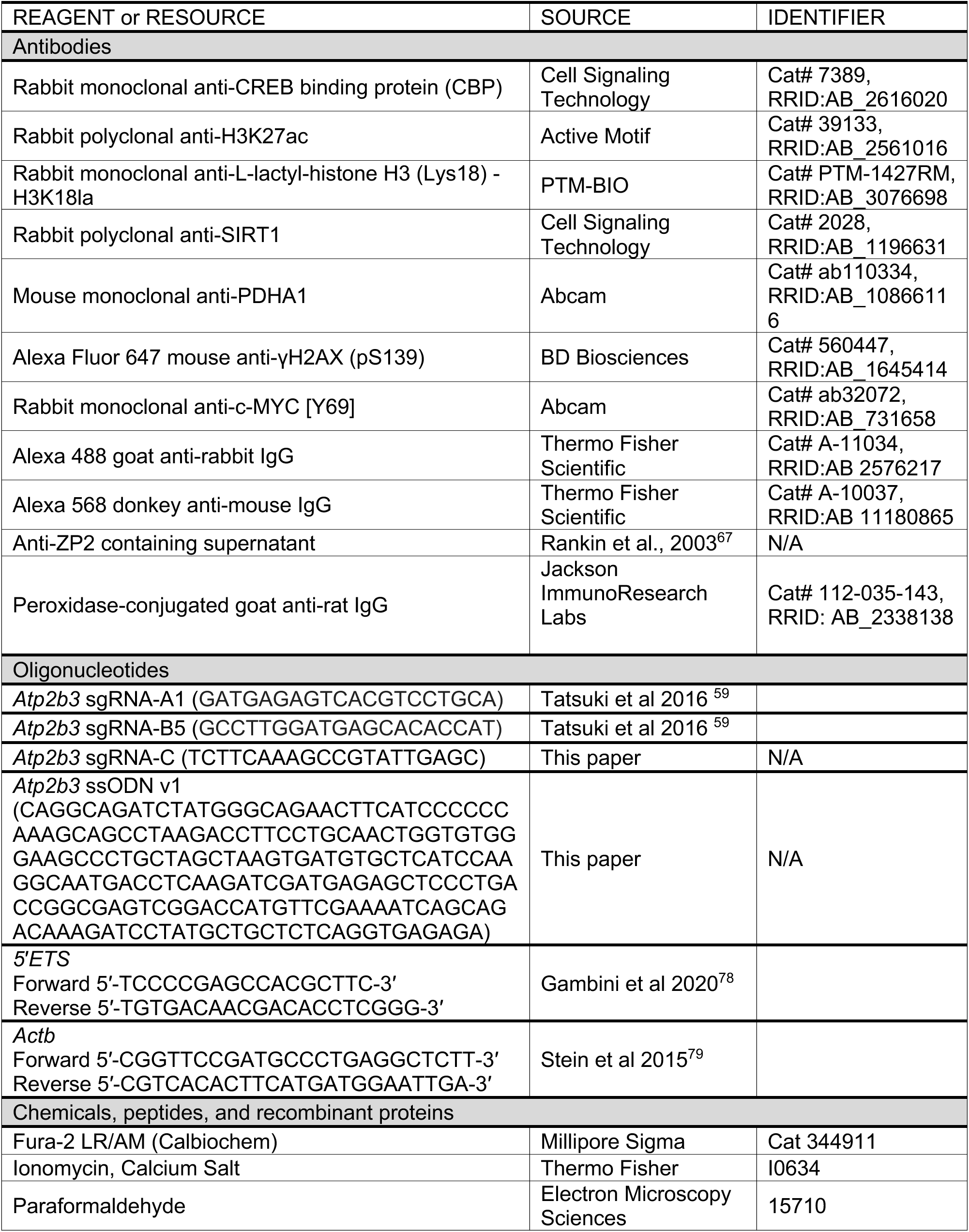
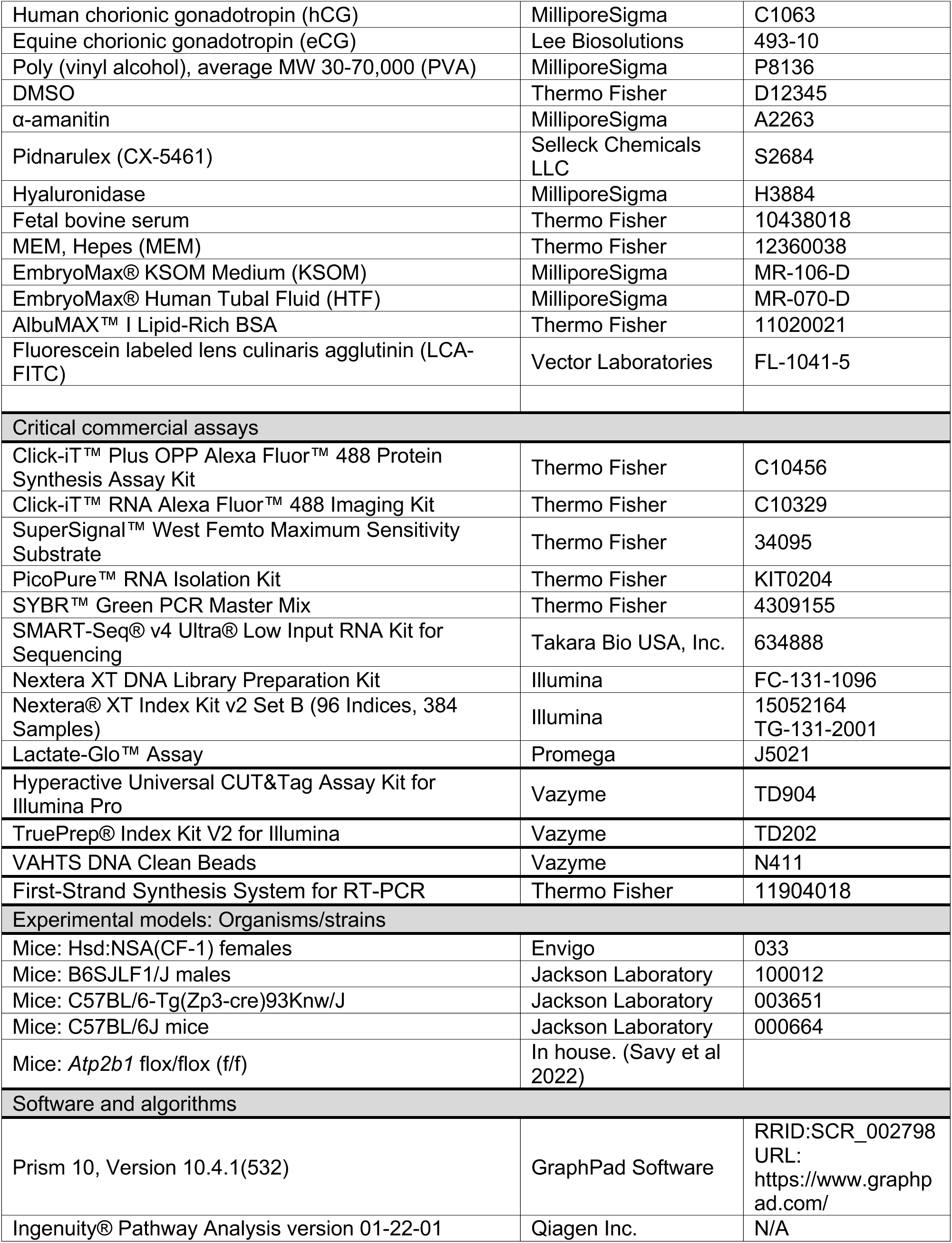
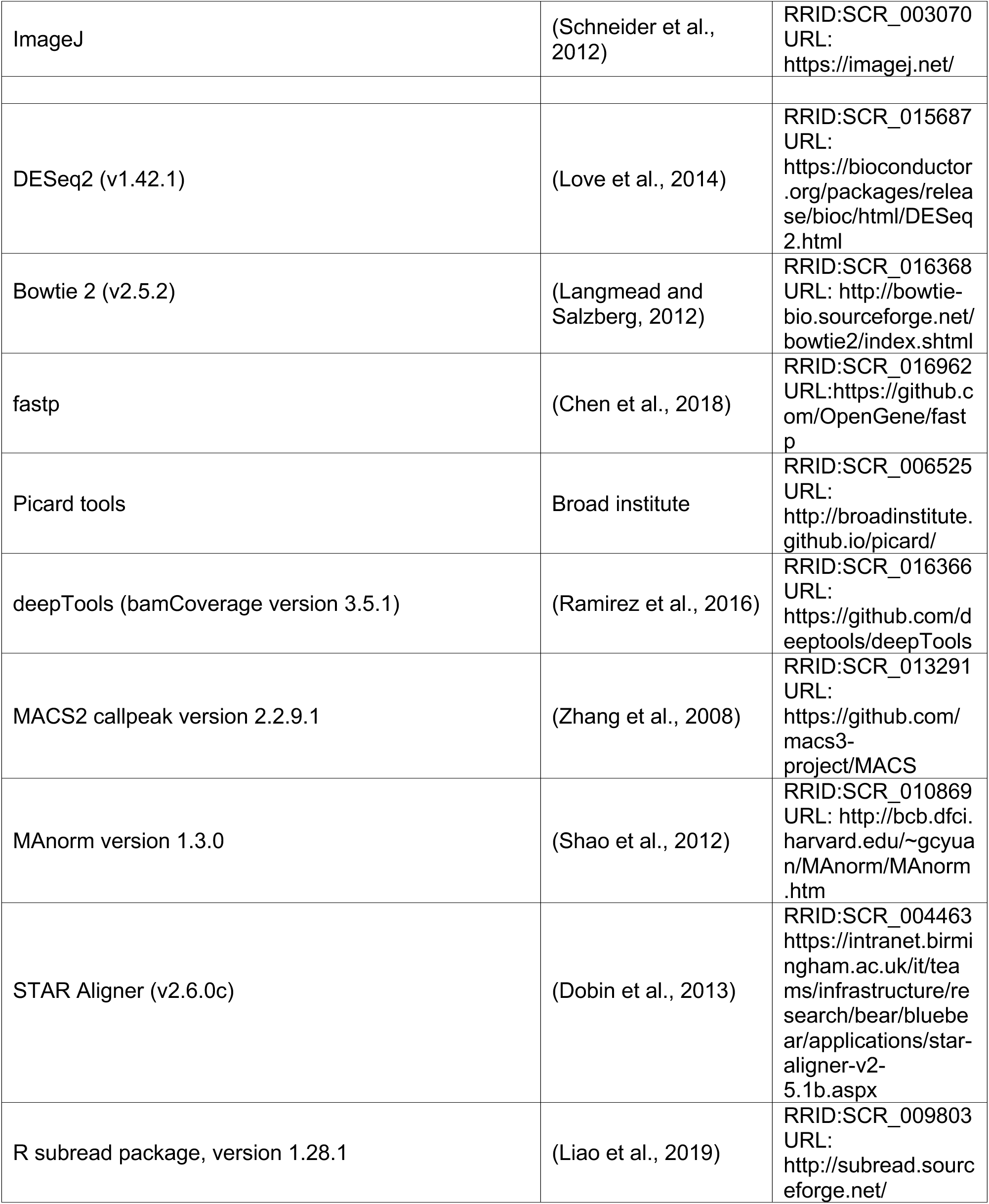

---

**Figure S1.**
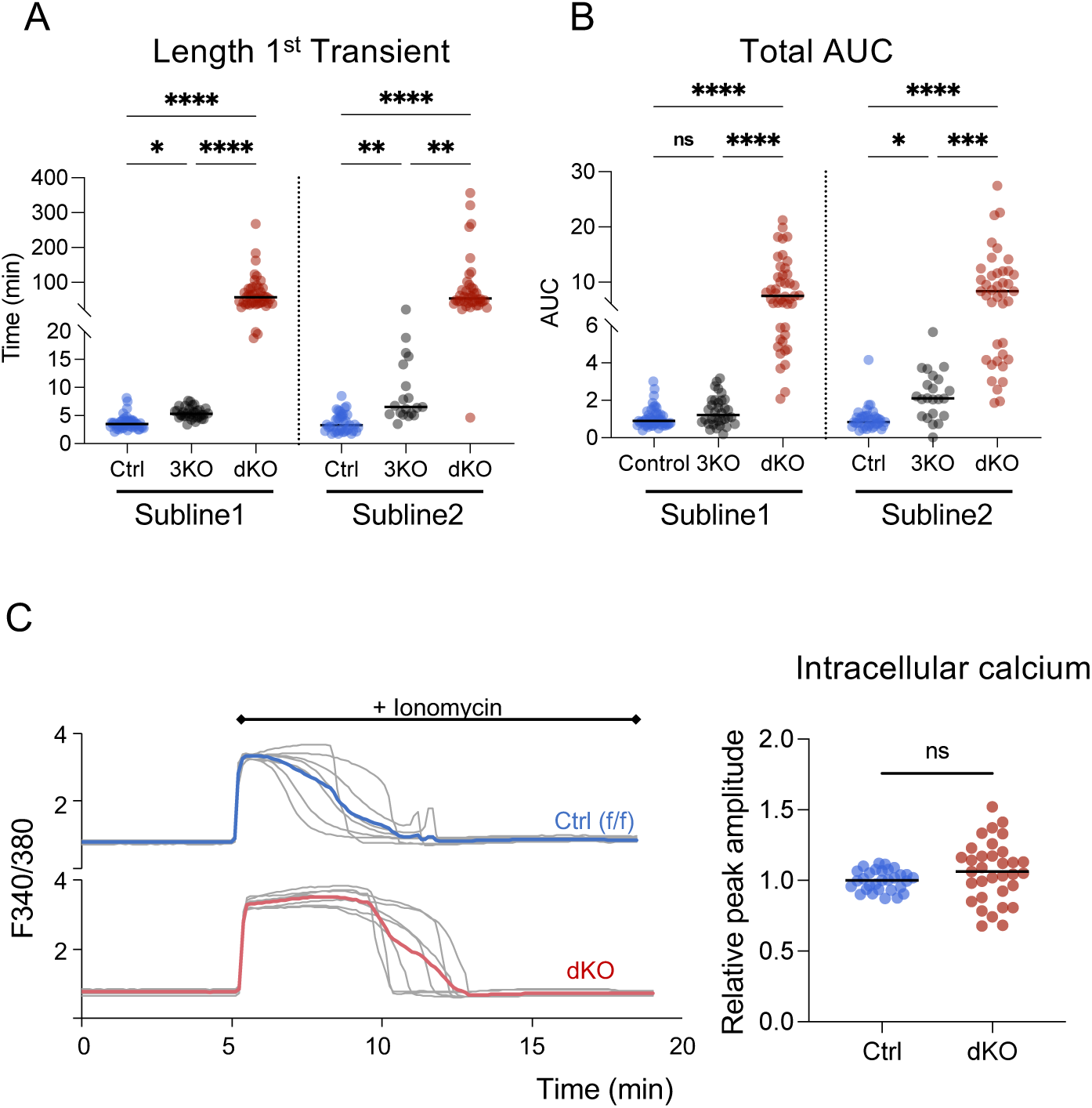
Analysis of ratiometric Ca^2+^ imaging during IVF of control (blue), PMCA3-KO (3KO, gray) and PMCA1/PMCA3 double KO (dKO, red) eggs. N = 3 independent experiments using 2 independent mouse sublines (Subline1, Subline2). (A) Length of the first Ca^2+^ transient. (B) Area under the curve (AUC) of calcium signal, relative to controls. For A-C, Kruskal-Wallis with Dunnett’s multiple comparisons test was performed; *p<0.05, **p<0.005, ***p<0.0005; ****p<0.0001; ns, not significant. (C) Ionomycin-induced Ca^2+^ release in eggs from control (blue) and dKO (red) females. Left, representative traces; colored lines are means of the experimental traces shown in gray. Right, peak level of Ca^2+^ released as an indirect indicator of Ca^2+^ stores. T-test; ns, not significant.

**Figure S2.**
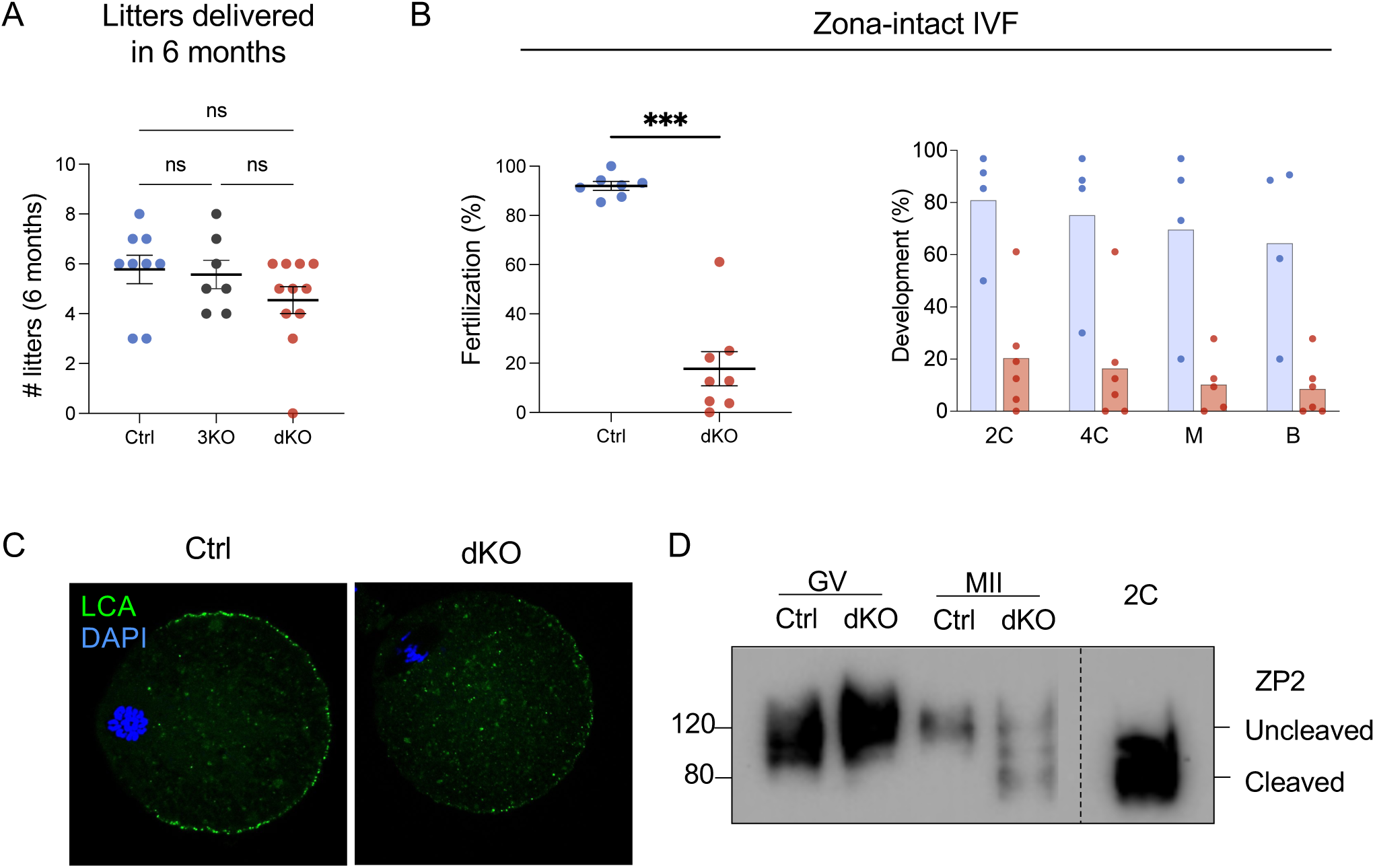
dKO eggs have premature zona pellucida hardening. (A) Number of litters born after mating Ctrl, 3KO, and dKO females to WT males; each dot represents the average litter size per female from 9, 7, and 11 breeding pairs, respectively, during a 6-month breeding trial. Kruskal-Wallis with Dunnett’s multiple comparisons test; ns, not significant. (B) Left, percentage of fertilized embryos following IVF with intact zona-pellucida. Each dot represents an independent biological replicate and horizontal bars indicate median. Mann Whitney test; ***p<0.0005. A total of 115 Ctrl and 170 dKO eggs were included in the analysis. Right, percentage of embryos that reached the various preimplantation embryo stages following IVF with intact zona-pellucida. Preimplantation embryo stages: 2C (2-cell), 4C (4-cell), M (morula) and B (blastocyst) stage embryos. (C) Representative images of LCA staining of cortical granules in zona-free MII-eggs from Ctrl and dKO females. (D) Representative immunoblot of glycoprotein ZP2 from Ctrl or dKO germinal vesicle stage oocytes (GV) or eggs (MII). Wild-type embryos at the 2C stage served as reference for ZP2 cleavage.

**Figure S3.**
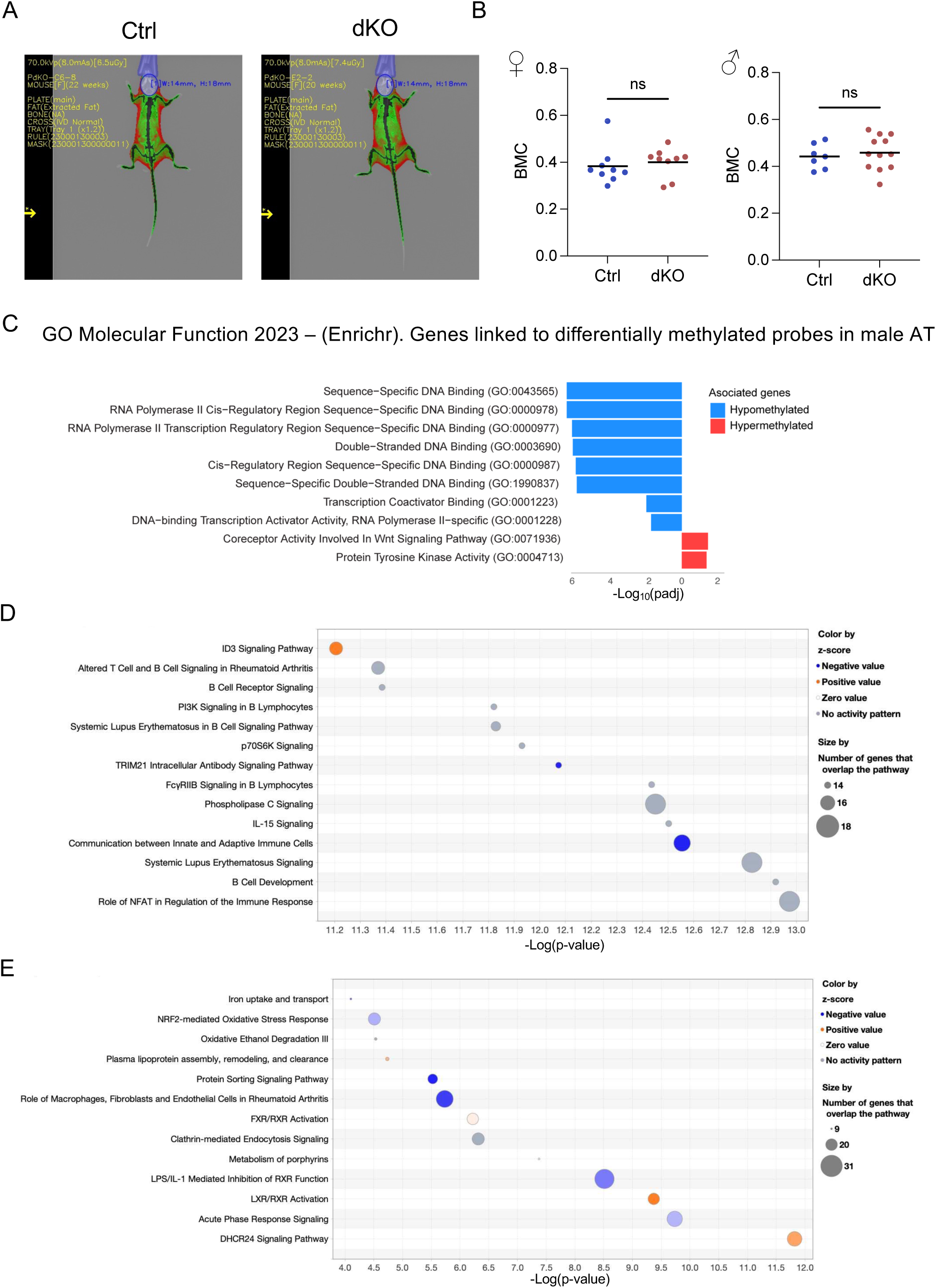
Increased Ca^2+^ at fertilization has long term effects on offspring health. (A) Representative images of body composition analysis for control (Ctrl, left) and dKO (right) derived offspring in adulthood. (B) Body mineral composition (BMC) analysis of Ctrl (blue) and dKO (red) derived offspring at adulthood. Each dot represents an individual animal. T-test; ns, not significant. (C) Top significant gene sets identified using Enrichr GO Molecular Function 2023. Gene Ontology analysis of genes associated with hypermethylated (red) or hypomethylated (blue) probes in adipose tissue from dKO males. (D-E) Top canonical pathways from Ingenuity Pathway Analysis of DEGs in adipose tissue from dKO vs control females (D) and males (E). Canonical pathways are displayed on the Y-axis and are sorted by p-value.

**Figure S4.**
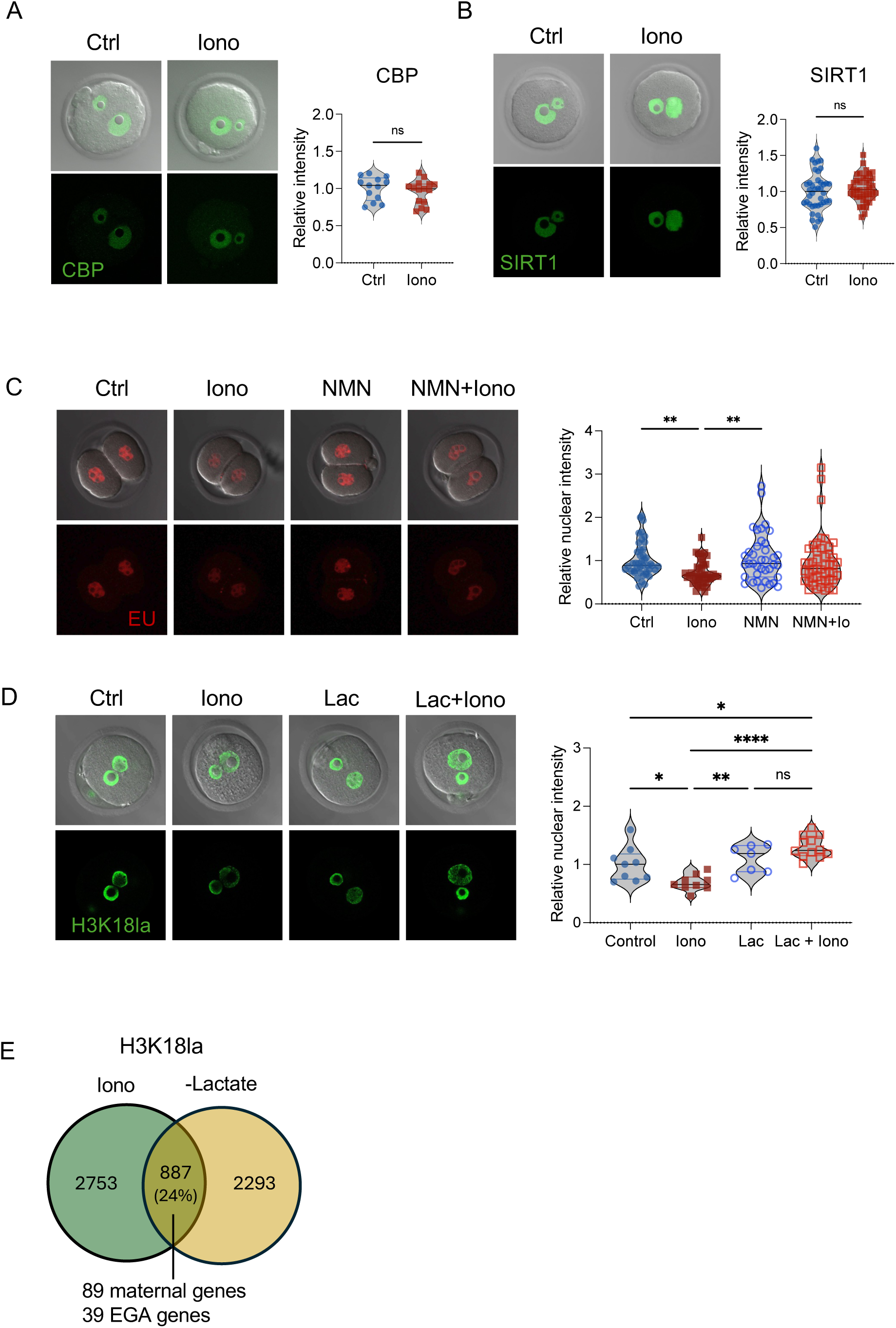
Exogenous lactyl-CoA but not NMN can mitigate Ca^2+^-mediated changes in reprogramming. (A-B) Representative images and quantification of nuclear staining of CREB-binding protein (CBP) and sirtuin 1 (SIRT1) in control (Ctrl) and ionomycin-treated embryos (Iono) at the 1C stage. Unpaired t-test; ns, not significant. (C) Representative images and quantification of global transcription activity (EU levels) in embryos at the 2C stage from Ctrl and Iono groups, as well as embryos cultured with exogenous nicotinamide mononucleotide (NMN) and NMN plus ionomycin (NMN+Iono). Kruskal-Wallis with Dunnett’s multiple comparisons test; **p<0.005. (D) Representative images and quantification of H3K18la levels in embryos at the 1C stage from Ctrl and Iono groups, as well as embryos microinjected with lactyl-CoA (Lac) and lactyl-CoA plus ionomycin (Lac+Iono). One-way ANOVA with Tukey’s multiple comparisons test; *p<0.05, **p<0.005, ****p<0.0001; ns, not significant. (E) Venn diagram comparing differential H3K18la peaks between Ctrl and Iono embryos at the 2C stage, overlapped with previously published differential H3K18la peaks in embryos cultured with and without lactate (-Lactate) at the 2C stage^19^.

